# Linking selection to demography in experimental evolution of active death in a unicellular organism

**DOI:** 10.1101/2024.06.28.601143

**Authors:** Nathalie Zeballos, Oceane Rieu, Stanislas Fereol, Christelle Leung, Luis-Miguel Chevin

## Abstract

Deciphering how natural selection emerges from demographic differences among genotypes, and reciprocally how evolution affects population dynamics, is key to understanding population responses to environmental stress. This is especially true in non-trivial ecological scenarios, such as programmed cell death (PCD) in unicellular organisms, which can lead to massive population decline in response to stress. To understand how selection may operate on this trait, we exposed monocultures and mixtures of two closely related strains of the microalga *Dunalliela salina*, one of which induces PCD, to multiple cycles of hyper-osmotic shocks, and tracked demography and selection throughout. Population dynamics were consistent between mixtures and monocultures, suggesting that selection on PCD does not involve strong ecological interactions. The PCD-inducing strain was maintained throughout the experiment despite an initial decline, by a combination of fast population rebound following each decline, and density-dependent competition dynamics near the stationary phase that were independent of these initial population fluctuations. As result of PCD maintenance, population decline in response to environmental stress was not counter-selected in our experiment, but persisted over 13 cycles of salinity. Our results highlight how analysing the demographic underpinnings of fitness and competition can shed light on the mechanisms underlying selection and eco-evolutionary dynamics.

## Introduction

Natural selection is intrinsically a demographic process, arising from the differential growths of genotypes with different vital rates (survival and fecundity) underlying fitness [1,2]. However beyond this uncontroversial statement, deducing selection from differences in demographic properties of genotypes often proves a non-trivial effort [3,4], and several important questions need to be addressed in order to decipher how selection operates on a trait, either directly or indirectly.

First, to what extent can absolute fitness of genotypes measured in isolation be used to predict selection when these genotypes are in competition [5]? It is well understood that what matters for selection is relative fitness, the difference between the relative growth rates of different types in competition [1,6,7]. However, measuring relative fitness comes with the technical challenge of being able to distinguish the competing genotypes. This can be achieved by relying on natural phenotypic variation such as colour polymorphism [8,9]; by inserting phenotypic markers such as fluorescent proteins [10]; or by sequencing standing genetic variants [11–13], or designed ‘barcode’ sequences [14]. Beyond these technical limitations, focusing solely on relative fitness can make it difficult to decipher the causes and mechanisms of selection across environments and ecological contexts. More insight may be obtained by studying the demographic responses of genotypes in isolation (absolute fitness) first, and comparing them to their differential growth in competition (relative fitness). This approach is particularly useful when analysing eco-evolutionary scenarios where rapid evolution is likely to influence population growth, for instance during evolutionary rescue [15,16].

Closely linked to the relationship between relative and absolute fitness is the question: To what extent are evolutionary dynamics determined by differences in intrinsic performances of genotypes, as opposed to interactions among these genotypes? Relative fitnesses of competing genotypes can be well predicted from their absolute fitnesses in isolation when these genotypes do not interact ecologically in ways that lead to frequency-dependent selection [5,17]. This condition is less likely to be met whenever one of the genotypes excretes beneficial or detrimental substances in the medium [18], or benefits from arriving or starting to grow earlier than others (priority effect) [19], for instance. Therefore, contexts where ecological interactions are suspected to be important are less likely to show consistency between fitness measured in isolation versus in competition. Reciprocally, finding similar demographic dynamics in isolation versus competition can help exclude some forms of interactions between genotypes, and more broadly delineate plausible underlying ecological causes for observed eco-evolutionary dynamics.

Even without frequency-dependent selection, the dynamics of selection may differ drastically between phases of rapid growth, where low population densities mean that genotypes compete little for resources, and phases with dense populations, where what matters is the ability to efficiently exploit these resources. This type of reasoning underlies classical arguments about r-versus K-selection [20], which have rejuvenated in recent years through new theoretical developments and empirical evidence in the wild [21,22]. Whether differences in maximum growth rate or competitive ability matter most for selection should depend on the extent to which different genotypes are limited by similar resources and, in an experimental context, on the frequency at which resources are renewed.

These general questions about the links between demography and evolution become especially important in contexts where genotypes differ non-trivially in their demographic responses to environmental change. For instance, it has been recently shown that two strains of the halotolerant microalga *Dunalliela salina* have drastically different demographic strategies in response to hyper-osmotic stress [23,24]. One of the strains undergoes programmed cell death (hereafter PCD), an active form of cell death that has been reported across a broad range of unicellular organisms in response to a variety of environmental triggers [25–27]. In *Dunalliela salina*, the strain inducing PCD undergoes a massive and rapid population decline at the onset of osmotic shock, while a closely related strain does not [23,24]. Strikingly, the declining strain later experiences a rebound phase characterized by fast growth in the same stressful environment, such that its monocultures may reach similar or higher population density than those of the non-declining strain after a few generations (about a week) [24]. If these dynamics in monoculture are good predictors of the outcome of competition between these strains, then we expect the genotype inducing PCD to be favoured by natural selection over successive osmotic shocks, and thus increase in frequency. This would in turn cause the (somewhat paradoxical) evolution of sharper decline in response to hyper-osmotic stress at the population level.

On the other hand, these predictions would be violated if ecological interactions perturb the outcome of competition. For instance, the non-declining strain could exhaust resources before the declining strain has recovered from its initial decline, preventing it from rebounding. This is an example of a priority effect [19], which could lead to competitive exclusion of the declining strain. Alternatively, dying cells from the declining strain may release resources or cues in the medium that fundamentally modify the growth rate of the co-occurring non-declining strain [28,29]. Even regardless of these ecological considerations, for the declining PCD strain to outgrow its competitor despite having initially declined, it needs to grow faster for sufficiently long, before reaching carrying capacity where growth is no longer possible. Furthermore, the sensitivity to competition near carrying capacity may differ between strains, thus influencing the outcome of their competition, and thus selection.

To investigate how these processes drive eco-evolutionary dynamics in our system, and better understand the – so far largely misunderstood [30] - putative benefit of PCD, we exposed monocultures and mixtures of clones of the declining and non-declining strains of *D. salina* to 13 cycles of hyper-osmotic stress (∼180 generations), and tracked demography and selection throughout. Our results reveal that the long-term maintenance of PCD can be explained by a complex interplay between decline-rebound dynamics and density-dependent competition near carrying capacity, emphasizing the importance of deciphering the demographic underpinnings of selection in experimental evolution.

## Materials & Methods

### Experimental evolution

We focused on two closely related *Dunalliela salina* strains that were previously shown to differ in their responses to a salinity rise [23,24]. The first one, CCAP 19/12 (hereafter strain A), sharply declines shortly after a salinity rise. This abrupt and massive decline (with up to 70% of the population disappearing in one hour), attributable to programmed cell death (PCD, see below), is then followed by a rapid rebound, characterized by fast population growth [23,24]. In contrast, the other strain CCAP 19/15 (strain C) does not exhibit any decline-rebound pattern, instead growing at a relatively constant rate. Interestingly, these previous investigations suggested that monocultures of strain A could reach similar or even higher density than strain C at the end of the rebound phase [24]. We therefore wanted to test whether this is also true in competition, and how this influences experimental evolution of PCD. Both strains were obtained from the Culture Collection of Algae and Protozoan (CCAP) in 2017, and were originally isolated from the same sampling site (North Sinai, Israel in 1976). Each of the 10 biological replicates per strain used in this study was founded from a single haploid cell in May 2019 [31], ensuring nearly isogenic populations per replicate (but with potential genetic variation among these clones).

To investigate competitive fitness among strains and the resulting experimental evolution of PCD, we exposed several mixtures of these strains to successive osmotic shocks, and tracked their population dynamics through time. Cell cultures were performed in 50 ml flasks (CELLSTAR®; VWR 392-0016), using custom-made artificial saline water complemented with NaCl [24,31]. Growth chambers were set to 12h:12h light:dark cycles, with light intensity at 200 μmol.m^-2^.s^-1^, temperature at 24°C, and randomized position. Mixed populations of *D. salina* were initiated with 50% of strain A and 50% of strain C, starting at a density of 50,000 cells/ml, and using a different biological replicate (i.e. isogenic population) for each mixture. These mixed populations were then exposed to 13 cycles of salinity changes, leading to ∼180 generations of experimental evolution, assuming a generation time of ∼1 day [32]. Each cycle consisted of a step at 2.4M NaCl (referred to as low salinity; but note that *D. salina* is a halophile, so this salinity is 4 to 5 times higher than seawater), followed by a hyper-osmotic stress where cells were subjected to a sharp salinity rise to 4.0M NaCl (high salinity). Cycles lasted 14 days, and we contrasted two temporal treatments: 4 days at low salinity followed by 10 days at high salinity (hereafter 4-10d cycle), or 7 days at each salinity (hereafter 7-7d cycle). We hypothesized that longer time at high salinity may provide the PCD strain more opportunity to recover from its initial decline when salinity is still high. At each transfer (i.e. salinity change), populations were diluted at a rate of 1/10, and the target salinity was achieved by mixing the required volumes of hypo-([NaCl] = 0 M) and hyper-([NaCl] = 4.8 M) saline media, after accounting for dilution of the previous salinity [31]. At the end of the 13 cycles of the assay, we genetically checked for the presence of both strains within each long-term mixed population (see Material S1).

In addition to these long-term mixed populations, our experiment included control lines subjected to the same salinity treatments, but restricted to the first 5 cycles (∼70 generations). These consisted first of monocultures with only strain A or C, with each replicate consisting of one of the isogenic lines used in the mixed populations. These monocultures not only served at controls for putative variation of the decline intensity over cycles (as observed in [24]), but are also necessary for estimating relative frequencies in the mixtures through the method described below. Another control consisted of mixing anew 50%-50% of strains A and C from long-term monocultures at each salinity rise. This allowed assessing whether the population dynamics of long-term mixed populations differed from those of populations with equal proportions of the two strains, thus quantifying the influence of evolution (frequency change) on population dynamics.

#### Population density measures

Since our interest is in demographic dynamics following each hyper-osmotic shock, we measured population density daily (except days 5 and 6) in the high salinity steps for the first five cycles (Fig. 1A) and the two last cycles (measurements for cycles 6 to 11 are explained in Material S2). In contrast in the low salinity steps, we only measured densities 1 hour after the salinity transfer to check the dilution rate, and at the end of the step (i.e. on day 4 or day 7 depending on the temporal treatment) to predict the initial density of long-term populations in the following high salinity step (if no decline occurred). This latter measure is also necessary to calculate the culture volumes required to start the control mixtures at 50,000 cells/ml at the next salinity transfer.

**Figure 1.**
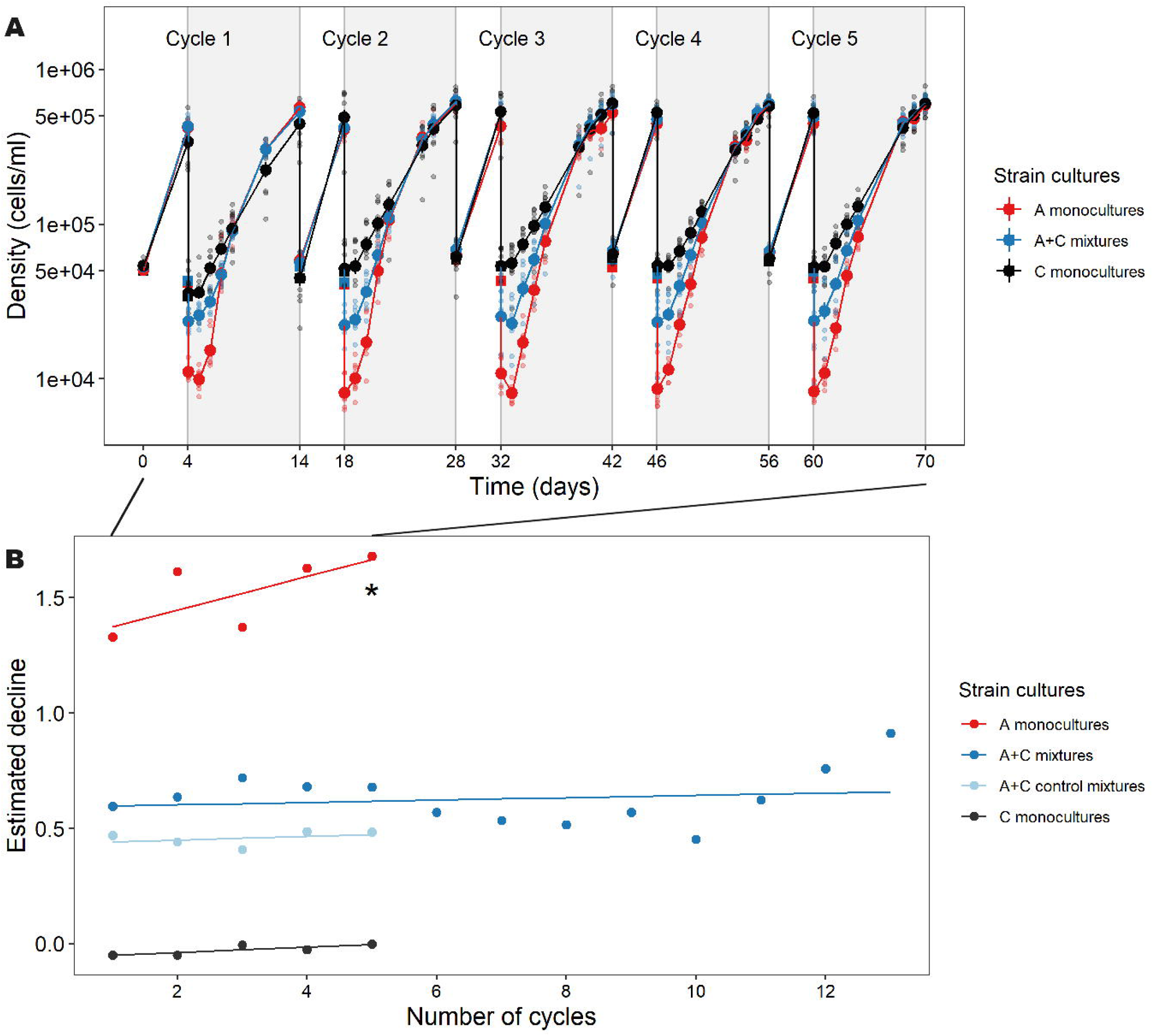
Demographic responses to successive salinity rises (4-10d fluctuation cycle). (A) Populations densities over the first five salinity cycles are shown for monocultures of strain A (red) and strain C (black), and long-term mixtures of A and C (blue). Small dots represent biological replicates (specific isogenic lines of each strain, and mixtures thereof), while large dots represent means over replicates (with SE not visible). Grey backgrounds correspond to the high salinity steps in a given cycle. (B) The estimated rate of initial decline one hour after the hyper-osmotic shock is shown over cycles, based on GLMs where the number of cycles was treated either as factor (dots), or as a continuous variable (lines). The star denotes p-value <0.1 for the slope of decline rate against cycle number.

We measured population densities by passing 200 μl samples of each population through a Guava® EasyCyte™ HT cytometer (Luminex Corporation, Texas, USA), with a laser emitting at 488 nm. *Dunaliella* cells emits natural fluorescence in red (695/50nm) and yellow (583/26nm) due to their chlorophyll α, allowing discrimination of live *Dunaliella* cells from other particles, as described in [24,31]. A decrease in population density between the time of transfer to the new high salinity environment and 1h after the transfer was assumed to be entirely due to PCD, as this population decline was cancelled in presence of a PCD inhibitor (see below and [24]). Furthermore, non-PCD dying cells could be discriminated based on their smaller size and weaker red and yellow fluorescence (see Fig. S1 and further details in [24]), but their number cannot account for the massive decline observed.

#### Caspase-3-like inhibitor assay

It has been shown that the intensity of decline in strain A can be diminished by an inhibitor of caspase-3-like enzymes [24], known to be a key player in the metabolic cascade leading to cell death [25,33–35]. We therefore investigated how this PCD inhibitor influenced strain frequencies and population dynamics in mixed populations. Both strains were first acclimated for 4 days at 2.4M. We then prepared 10 mixtures (50-50%) at *N*_0_ = 50,000 cells/ml, and 15 monocultures of each strain at two initial densities: 10 at *N*_0_ = 50,000 and 5 at *N*_0_ = 25,000 cells/ml, respectively matching the total density of mixed populations and of each individual strain therein, in total culture volume of 10ml. We applied 2 inhibitor conditions (with or without PCD inhibitor), reaching a total of 80 flasks. In the inhibitor treatment, a total number of 500,000 cells per flask were treated with 10 μM Z-VAD(Ome)-FMK (CellSignaling) caspase inhibitor, and incubated in the dark for 30min at 24°C, before being subjected to a hyper-osmotic shock at 4M NaCl. Population densities were measured 30 min and 2 hours after the hyper-osmotic transfer, and then daily tracked over 10 days. Since the inhibitor is supplied as 1 mg of powder and has been diluted in 213.9 μl of DMSO with membrane penetration action [36], we also checked that this solvent did not prevent decline of strain A monocultures (see Material S3).

### Demographic and competition analysis

#### Decline intensity

To quantify the intensity of population decline following each transfer to high salinity, we used a GLM with cytometer cell count one hour after the transfer (in day 0) as response variable, a log link function, negative binomial error structure, and log expected initial population density (based on pre-transfer density and the dilution rate) as offset (following [23,24,31]). We extracted the decline rate from those GLM as minus the linear predictor of the GLM (hence, on the log scale), such that steeper decline is represented by a larger positive value [24]. We then computed population size reduction in % from the estimates of the GLMs. The fixed effects were the type of culture (monoculture or mixture), temporal treatment (4-10 vs 7-7 days), and the number of cycles (either as categorical or as continuous variable). The latter effect was used to investigate whether the intensity of decline varied over cycles of salinity fluctuations. In mixed populations (with both A and C strains), such a change in decline rate over cycles could result from a change in frequency of the declining strain A over cycles, indicative of selection. However, it could also result from a change in the decline rate of strain A over cycles. This can be investigated by tracking putative changes in the decline of strain A monocultures over cycles.

#### Competition analysis

In most mixed populations, the two strains differed in their natural yellow and red fluorescence, despite some overlap (Fig. S1). We thus designed a mixture analysis method to estimate the relative frequencies of both strains, and from this their absolute population densities in the mixture, accounting for variable uncertainty among mixture types (clones used for each strain), as detailed in Material S4. We validated this method using mixed populations where we controlled the relative frequency of strain A, either experimentally or virtually (Material S6 and Fig. S2). This framework for estimating frequency in a mixture using monocultures as reference is broadly applicable in any competition experiments with genotypes/species distinguished by their phenotypic traits, beyond the specific context we investigate here.

We then computed the *per-capita* growth rate per day of each strain in mixed populations during the exponential phase over 1000 simulated data sets, to account for uncertainty in the estimator of frequency of strain A. Finally, we fitted linear models (LM) and generalised additive models (GAM) with the *per-capita* growth rate as response variable, and the densities *N̂*_*t−τ*_ of both strains or the total densities in mixtures as fixed effects (see Table S1 for details), on each of the 1000 simulated dataset, and then obtained combined estimates with standard error using Rubin’s rules (see Material S5).

#### Exponential growth rate in the inhibitor assay

We applied a generalised linear model on population densities (from day 2 to 5) of strain A monocultures in the PCD inhibitor assay, to estimate the growth rate during the exponential phase as in [24]. Here, time was included as a continuous explanatory variable, estimating the rate of exponential growth (linear trend on log scale), and interactions of time with the other fixed effects (i.e. inhibitor treatment and initial density) estimated effects of these factors on the maximal exponential growth. All statistical analyses were performed on Rstudio (R version 4.2.0) using packages MASS (version 7.3.56) [37], stats (version 4.2.3), mvtnorm (version 1.1.3), mgcv (version 1.8.33), mice (version 3.14) [38].

## Results

### Population decline occurs repeatedly over successive hyper-osmotic shocks

The population dynamics over the first 5 cycles of salinity are shown in Fig. 1. Soon after each hyper-osmotic transfer, monocultures of strain A first declined sharply (population size reduction averaged over 7-7d and 4-10d treatments and over 5 cycles: 72.5%, SD= 6.8%), before rebounding through fast growth, eventually reaching a similar density as monocultures of the non-declining strain C, consistent with results in [24]. The decline-rebound pattern of strain A was repeated over successive hyper-osmotic shocks, with a decline intensity that slightly increased over cycles (number of salinity cycles used as a continuous variable, Table S2, P = 0.044, red line in Fig. 1B). The same pattern was found regardless of cycle treatment (Fig. 1A & S3A), but the increase in decline intensity over cycles was not significant in the 7-7d treatment (Table S2, P = 0.914). The persistence of decline intensity over cycles indicates that this response is not a mere passive consequence of the elimination of cells in bad condition [24].

To investigate whether the declining strain could be maintained in competition with a non-declining one, we exposed mixed populations comprising both strains to the same succession of cycles. Based on the results of monocultures, we may predict that the declining strain A should outcompete the non-declining strain C, leading to a more marked decline over cycles in mixed population. However, more complex dynamics may play out in mixtures, for instance if strain C exhausts resources before A is able to rebound (priority effect [19]). Here mixed populations maintained a similar rate of decline (population size reduction averaged over 7-7d and 4-10d treatments and over the 13 cycles: 45.6%, SD= 6.5%), intermediate between those of strain A and C monocultures (population size reduction averaged over 7-7d and 4-10d treatments and over 5 cycles: -4.3%, SD = 4.0%), across hyperosmotic shocks (Fig. 1B, Table S2, P = 0.913), suggesting that no strain was able to fully outcompete the other after 180 days.

### Strain frequencies fluctuate within a cycle but are stable over cycles

To further understand how these demographic dynamics relate to selection, we estimated strain frequencies in the mixed populations by taking advantage of their phenotypic differences (as measured by flow cytometry), and using the monocultures from the same day as references (Fig. S1). Strain A frequency strongly fluctuated in the days following hyper-osmotic shock. It was pretty low shortly after the osmotic shock (averaged over 5 first cycles for the 4-10d treatment: 0.36, SD = 0.17), before increasing until it became dominant in the mixture at the end of the high salinity step (averaged over 5 first cycles for the 4-10d treatment: 0.64, SD = 0.13, Fig. 2), in almost every cycle (Figs. 2 & S4). This pattern is consistent with what could be predicted from the decline-rebound pattern in population density observed in monocultures. In the control mixtures, in which the frequency of strain A was set to ∼50% before each hyper-osmotic shock, this frequency started at lower value than in long-term mixtures, when first measured in the hypersaline environment (averaged for for the 4-10d treatment: 0.11, SD = 0.04). This suggests that the proportion of strain A before the hyper-osmotic shock (in the previous low salinity step) was greater than 50% in the long-term mixtures. We confirmed the stable coexistence of both strains in all long-term mixtures by genetically detecting both strains after 13 salinity cycles (Fig. S5).

**Figure 2.**
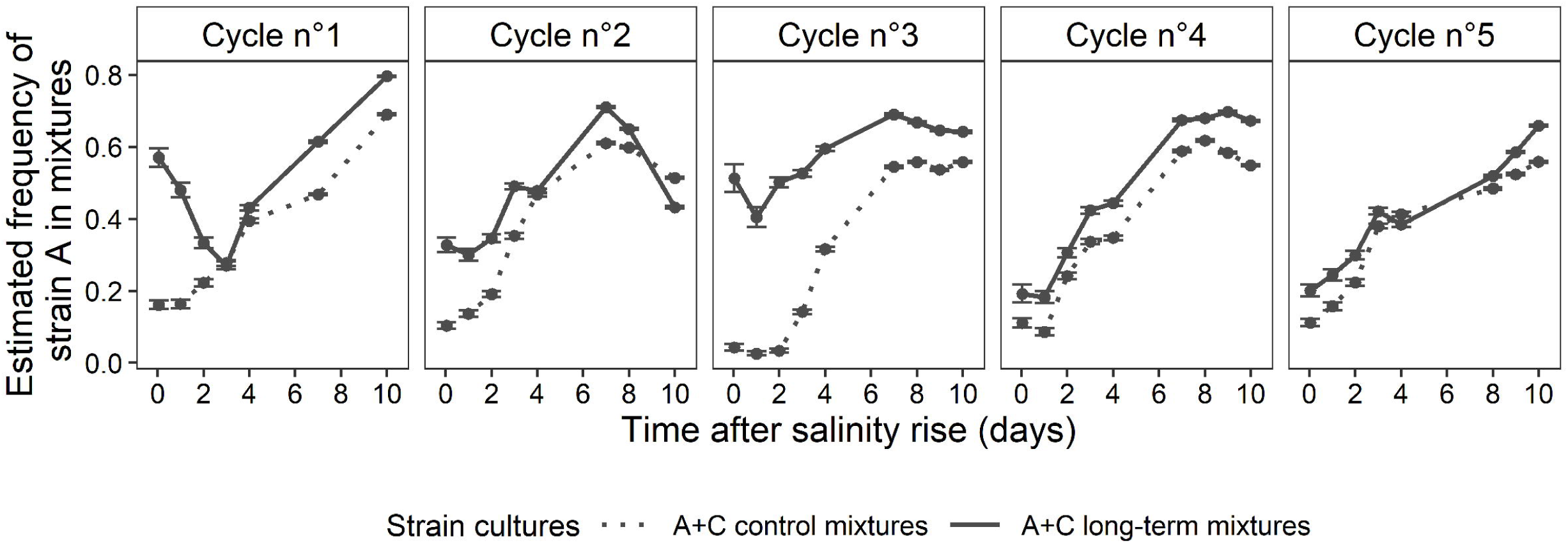
Frequency of declining strain over 5 successive salinity rises (4-10d fluctuation cycle). The proportion of strain A in mixed populations, as estimated from the cytometric traits of cells, is shown against time for long-term mixtures (solid lines), and control mixtures (dotted lines) that were freshly composed by mixing 50-50% of each strain before each salinity rise. Means and 95% confidence intervals were obtained on the logit scale by inverse-variance weighting over the 10 population lines, to account for the variable uncertainty of estimates among biological replicates (isogenic lines of each strain), and back-transformed to the arithmetic scale using the delta method.

Since strain A grows faster than strain C during its rebound phase (Fig. 1A; also [24]), we hypothesized that longer hypersaline phases would leave more time for A to outgrow C in mixed populations. However, the dynamics of strain A frequency in the 7-7d cycles was similar to that in 4-10d cycles, over the first 5 cycles (Fig. S3B, initial frequency 0.38, SD = 0.10) as well as in the last two cycles (Fig. S4). This suggests that 7 days were sufficient for strain A to reach high enough frequency at the end of each hypersaline phase (averaged over 5 first cycles for the 7-7d treatment: 0.68, SD = 0.09) to not be outcompeted at the next hyper-osmotic shock. This is also consistent with the stable frequency observed between days 7 and 10 under 4-10d cycles (Fig. 2), suggesting that differences in growth rates of both strains were no longer significant drivers of selection after 7 days.

### To what extent does demography explain selection?

To further elucidate how these selective dynamics of changes in relative genotype frequencies emerge from demography, we calculated the density of each strain in the mixed populations from their estimated relative frequency (Material S4). The demographic dynamics of each strain in mixtures (Fig. 3, dashed lines) was qualitatively consistent with that in monocultures (Fig. 3, solid lines). In the mixtures, strain A had lower density than strain C just after the hyper-osmotic shock as a consequence of PCD (averaged over the 5 cycles for the 4-10d treatment: 8,867 cells/ml, SD = 4,368 cells/ml vs 15,778 cells/ml, SD = 4,541 cells/ml), but its higher exponential growth rate during the rebound allowed it to reach higher density at the end of the hypersaline phase (averaged over the 5 cycles for the 4-10d treatment x1.7: 3.8×10^5^, SD = 6.2×10^4^ vs 2.2×10^5^, SD = 8.8 x10^4^ in Fig. 3; see also Fig. S4C). This growth advantage during rebound was more pronounced than expected based on monocultures: in Fig. 3, the dashed red line crosses the dashed black line earlier, and their final difference is larger, than for continuous lines (monocultures). This probably occurs because the density of each strain starts lower in mixtures than in monocultures (since they only compose half of the population), allowing growth differences to accrue over a longer exponential phase (Fig. 3, Fig. S4C). However, the outcome of this process should also depend on how density dependence acts within and between strains.

**Figure 3.**
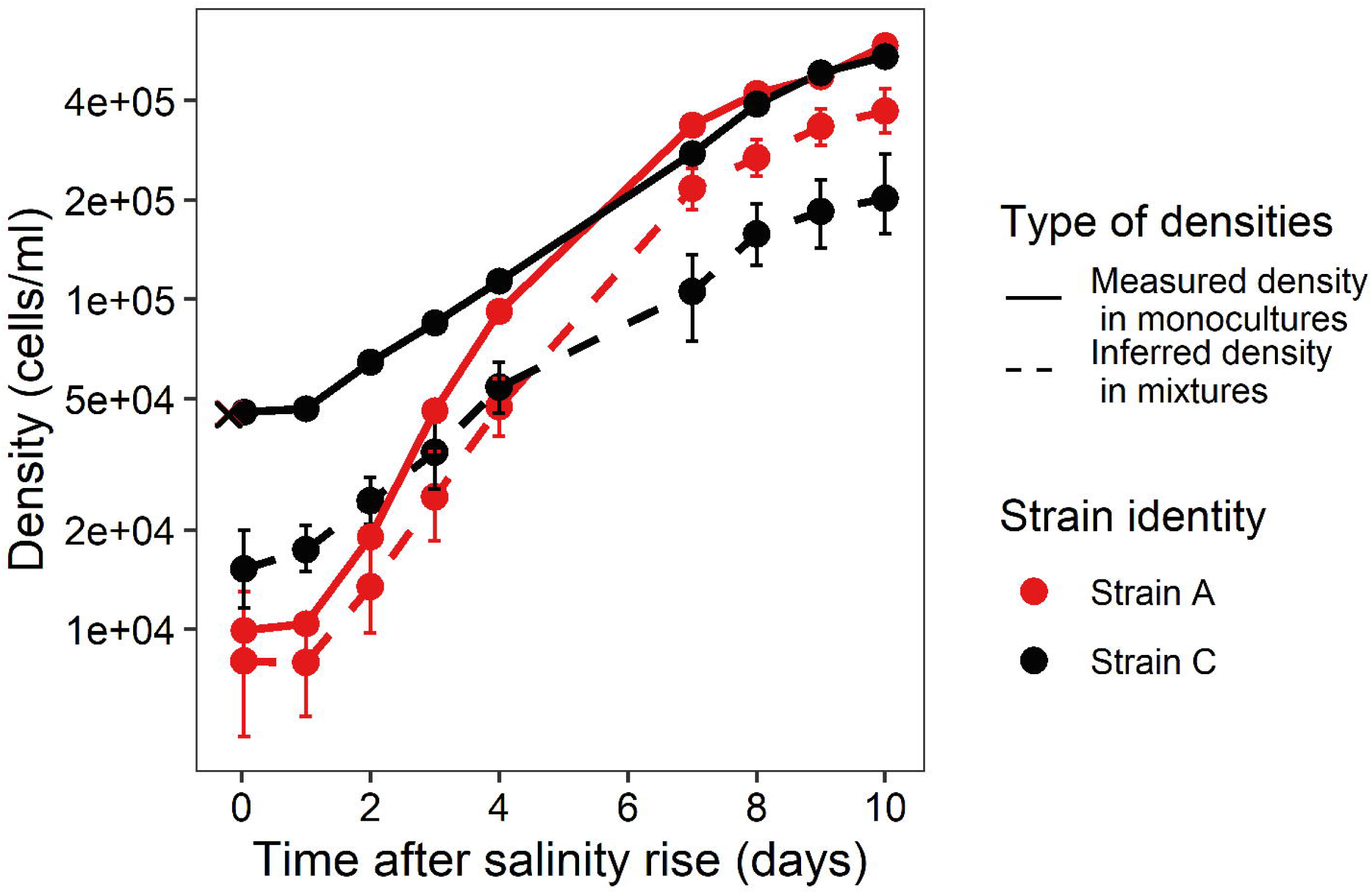
Growth rates in monocultures versus mixtures. Mean population densities across replicates in monocultures (solid lines; direct measurements), and long-term mixtures (dashed lines; inferred from the estimate of relative frequencies), averaged over the first 5 high salinity steps, are shown for the 4-10d fluctuation cycle. For the monocultures, we computed the mean and standard error (not visible) over 10 isogenic lines. For the mixtures, we used inverse-variance-weighted mean (eq. S9, computed with the delta method) and 95% confidence interval, to account for variable precision in estimation of strain A frequency among biological replicates. The crosses at time 0 (overlapping for red and black) correspond to the expected initial densities for monocultures, based on the known density before the 10% dilution.

To better understand how the growth of each strain is impacted by both intra and inter-genotype competition in the mixtures, we calculated the sensitivity of their *per-capita* growth rates to population density (Fig. 4, Table S3 for LM 1, see Fig. S6 for 7-7d treatment). We observed a significant negative effect of self-density on the *per-capita* growth rate of each strain (P = 5.34e-08 and P = 2.48e-06, Table S3), a classic signature of negative density dependence (i.e., regulation of population growth). The model that included densities of both self and the competitor strain as predictors (LM 1) showed that both strains are more sensitive to their own density than to that of their competitor, but that this is more pronounced for strain A (x4.0 for *r*_*A*_ vs x2.5 for *r*_*C*_, Table S3). Somewhat surprisingly, the linear model where the growth of strain A depends on the total density (LM 2, Table S4) performed poorly (higher AIC for LM 2 in Table S1), contrary to the expectation that for closely related strains, competition should depend on the total number of individuals, regardless their genotype.

**Figure 4.**
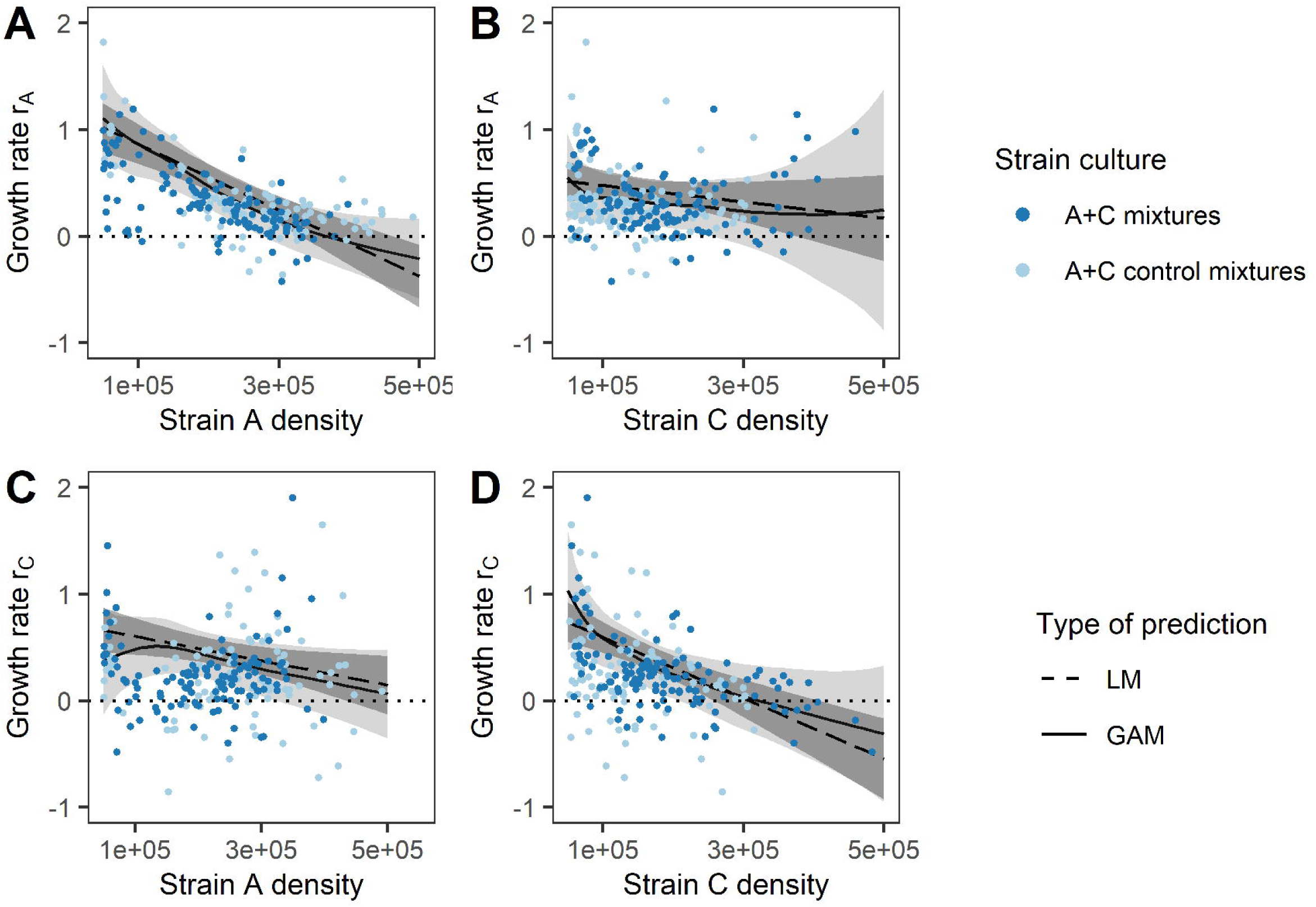
Intra- and inter-strain competition. The *per-capita* growth rate of strain A (A-B) or strain C (C-D) is shown against the density of strain A (A, C) or strain C (B, D), in the 4-10d fluctuation cycle. Each point corresponds to the *per-capita* growth rate over the interval between two subsequent population measurements, plotted against density at the first measurement. The predictions from linear models (LM1, straight dashed lines and dark gray 95% CI) and generalised additive models (GAM1, continuous lines and light gray 95% CI) are also shown, pooling estimates from 1000 resimulated datasets to account for uncertainty in *r* and *N*, and setting the density of the strain not represented in the x-axis to its median. We set the y-axis maximum to 2 for the sake of graphical clarity, which led to removing 2 dots in A-B (*r_A_* = 2.3; 3.9) and 1 in C-D (*r_C_* =2.4).

The fitted linear model (LM1) corresponds to the classic Lotka-Volterra model of competition, where the *per-capita* growth rate of the strain A is defined by

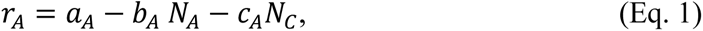

with *a*_*A*_ the intercept of the linear model (growth rate of strain A when both strains are at low density), and *b*_*A*_ and *C*_*A*_ the slopes of density dependence relative to self and the competitor, respectively (Table S3) (exchanging subscripts A and C yields the reciprocal equation for strain C.) In this model, the criterion for coexistence of two types is that each one is able to grow when at low density while the other is at demographic equilibrium [39]. In terms of parameters of the statistical model, this translates for the growth rate of strain A to

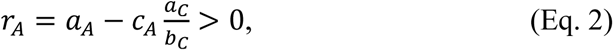

where 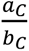 is the equilibrium population size of strain C in monoculture (satisfying *r*_*C*_ = *a*_*C*_ − *b*_*C*_ *N*_*C*_ = 0). The reciprocal invasion condition for strain C is 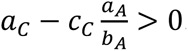. Here we found 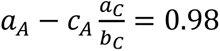 and 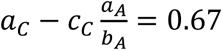, so the model predicts coexistence of both species. Furthermore, the equilibrium densities of both strains at the end of the stationary phase can be found by jointly solving for *r*_*A*_ = 0 and *r*_*C*_ = 0, yielding

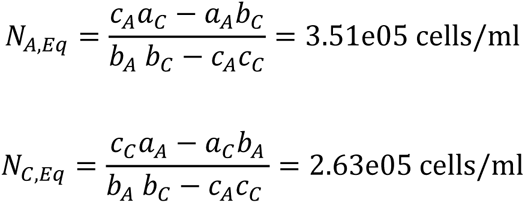

from the estimates in Table S3. From this, we expect the relative frequency of strain A at the end of the stationary phase to be 0.57, which is similar to the observed frequency at the end of each cycle (Fig. 2; average over 5 first cycles: 0.64). This shows that, although the rapid rebound that follows initial decline is critical to the maintenance of the PCD strain A, competition near the stationary phase is instrumental in stabilizing its frequency over cycles and promoting coexistence with the non-PCD strain C, largely independent of the initial decline-rebound dynamics. Similar conclusions were drawn using more flexible models (generalised additive models GAM, namely spline smoothers in the R package mgcv) that make fewer assumptions about the shape of density dependence (continuous lines in Fig. 4, Table S1).

### How does a PCD inhibitor affect competition in mixtures?

To demonstrate the role of programmed cell death (PCD) in these eco-evolutionary dynamics, we applied a caspase-3-like inhibitor to mixed populations composed of 50% of each strain, just before the underwent hyper-osmotic shock (Fig. S7). We estimated strain frequencies using monocultures as references, as explained above and in Material S4 (Figs. S7-S8), and then inferred the density of each strain in the mixtures (Fig. 5). While the treatment without inhibitor led to a sharp initial decline of strain A (estimated population size reduction in two hours of 73.4%) as previously, no population decline was observed for strain A subjected to PCD inhibitor (estimated population size reduction in two hours of -10.0%), confirming that the initial decline was attributable to a form of programmed cell death (as also shown in [24]). Interestingly, the mean growth rate of populations of strain A that underwent initial decline (without inhibitor) was higher than that of populations that did not decline (with inhibitor), over a similar range of densities (in mixtures, compare slopes of red vs dashed red lines between eg days 2 and 5 in Fig. 5; for monocultures see Table S5 P = 3.55e-11). In addition, the initial density of strain A monocultures without PCD inhibitor did not affect their growth rate during the exponential phase (Table S5, P = 0.451), consolidating the idea that the demographic rebound after the decline is not just a density-dependence effect due to relaxed competition. Eventually, strain A in mixtures reached a similar density with and without inhibitor (Fig. 5). The population dynamics of the non-PCD strain C were similar under both conditions, despite its competitor having markedly different responses, supporting the idea that the growth of strain C is little sensitive to presence of the other strain.

**Figure 5.**
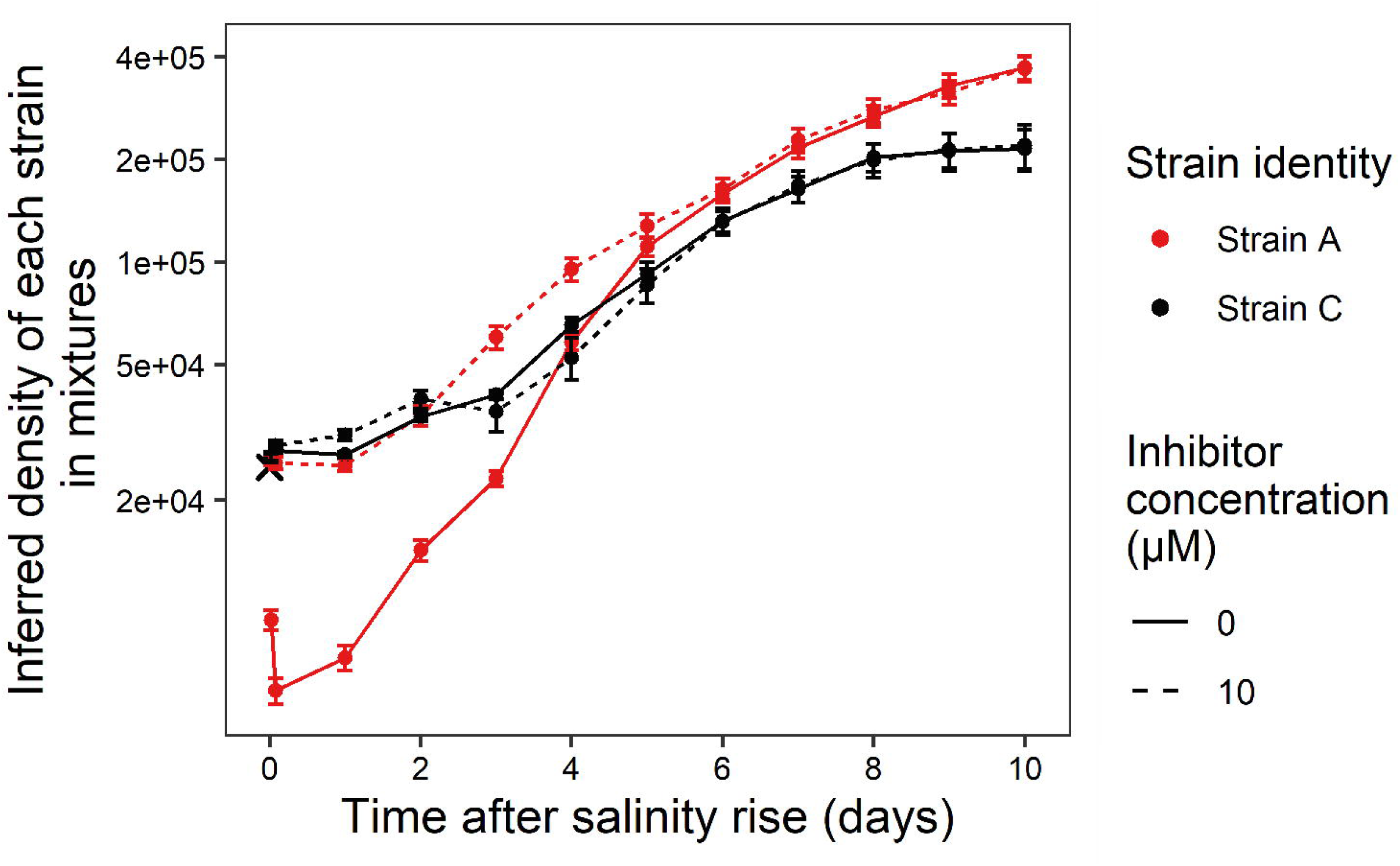
Impact of a programmed cell death inhibitor on growth in competition. Densities of each strain (colours) in the mixtures are shown under different doses of PCD inhibitor (line type), against days following a single salinity rise. Transfers occur at day 0, and the first measurements were made 30 min, and then 2 hours after the transfer. The cross corresponds to the expected initial density. Filled symbols are weighted averages (accounting for uncertainty in distinguishing the strains in mixtures) over 10 replicates, and error bars indicate confidence interval at 95.

## Discussion

We used a long-term competition experiment with two strains of the microalgae *Dunaliella salina*, one of which triggers programmed cell death (PCD) in response to hyper-osmotic shock, to investigate interconnected evolutionary ecology questions: (1) How can absolute fitness (population growth rate) of genotypes in isolation predict their relative fitness in competition, and thus the outcome of natural selection? (2) How does the outcome of this process depend on the demographic phase of population growth (exponential vs stationary), and what is the role of competition in long-term coexistence? (3) How can programmed cell death be maintained by selection, despite causing massive population decline? Our results provided important insights about all these questions, as we elaborate below.

### Relative and absolute fitness

Being able to measure relative fitness is crucial in experimental evolution, but is technically challenging as it requires being able to distinguish competing genotypes [10,11,14]. Here, we took advantage of the natural pigment fluorescence to discriminate the two strains in mixtures, and thus estimate their frequencies and relative fitnesses. Our method based on analysis of mixture distributions (Material S4) allowed us to estimate frequencies more precisely, and at much lower cost, than using strain-specific amplicon sequencing (as done in [11] for the same strains). This allowed us to increase temporal resolution (daily measurements), thus yielding more insights into the non-trivial eco-evolutionary dynamics in this system.

We have shown that the population dynamics in monocultures were broadly consistent with those of mixtures (Fig. 3), such that absolute fitness in monocultures partly predicted selection in mixtures. On the other hand, selection was strongly density-dependent, such that using exponential growth rates as proxies for fitness would not have been relevant to correctly estimate the net coefficient of selection over a cycle [5]. This was true regardless of the type of cycle (Fig. S6), and even though the transfer to the next salinity was achieved when strains had experienced less time in the hypersaline phase in the 7-7d cycle. Therefore in our experiment, a selection coefficient defined from differences in exponential growth rates, even averaged over the decline-rebound pattern for strain A, would have failed to predict the outcome of selection even within a cycle. Instead, the relative strain frequencies were largely driven in the long run by differences in their competitive abilities.

### Long-term coexistence of alternative strategies

Based on the competition-relatedness relationship, also known as the limiting similarity hypothesis [2,40–42], we might expect that the two strains we studied should exert strong competition on each other, since they are closely related. From this principle, we might have predicted exclusion of the declining strain A over cycles, since it starts growing later and at much lower density following each hyper-osmotic shock, such that strain C would benefit from a priority effect. Interestingly, we found that instead of responding to the total population density (Table S1, see AIC for LM 2 and GAM 2), as expected for closely related genotypes, the growth rates of both strains A and C were mostly sensitive to their own densities. This suggests that these strains have different ecological strategies in the same environment, as also indicated by their strikingly different demographies (Fig 1A, Fig 3). Such different ecological strategies between strains might arise from the use of different resources, such as light wavelength or depth in water column, or from different affinities to the same nutrient, among other possible explanations. In any case, this weak sensitivity to each other’s presence allowed their stable coexistence over 13 cycles of salinity changes (Fig. 1B and DNA confirmation Fig. S3), consistent with the fact that they have been sampled in 1976 from the same brackish site [43]. Previous work has shown that strain A is almost neutral against strain C at constant salinity 3.2M NaCl [11], but strongly counter-selected at constant 2.4M NaCl (our lower salinity here). Interestingly, although the dynamics of selection is here more complex because of the decline-rebound dynamics at each salinity rise, the long-term coexistence implies selective near-neutrality across cycles, for a mean salinity of 3.2M NaCl per cycle. Hence, fluctuating and constant salinities with a same mean 3.2M led to similar net selection, but probably for different reasons owing to their different demographic dynamics.

### Selection on programmed cell death

Beyond the general goal of understanding how selection emerges from demography, a major aim of our competition experiment was to determine whether, and how, selection can favour (or at least maintain) programmed cell death (PCD), a trait that seems detrimental since it leads to the death of individuals and the decline of populations. The distinctive decline-rebound pattern of strain A in response to osmotic shock was entirely removed by exposure to an inhibitor of caspase3-like activity (Fig. 5), confirming that it was a form of active death, or PCD (consistent with the results in [24]). A previous study with these strains has further shown that the intensities of decline and rebound were correlated across conditions, such that conditions more favourable to growth (more nutrients, more light) were associated to both steeper decline and greater rebound following hyperosmotic shock [24]. The ecological and/or physiological mechanisms behind the decline-rebound pattern, observed both in monocultures and in mixtures, remain to be elucidated. Three potential hypotheses were proposed in [24]: an altruistic release of substrate by the dying cells, heterogeneity in cell state, or a trade-off between halotolerance and reproduction. Further investigation of these hypotheses through modelling may help distinguish them, based on the dynamics they predict for both monocultures and mixtures.

Regardless of the mechanism driving natural selection on this trait, our results demonstrate that a PCD-inducing strain can be maintained in competition against a closely related non-PCD strain, over numerous successive osmotic shocks. This maintenance clearly relies on the rapid demographic rebound that follows the sharp decline due to PCD, allowing strain A to recover from its drastic initial demographic disadvantage and not be eliminated. On the other hand, the stable frequency of the PCD strain in the long run is mostly explained by competitive interactions at high density. These interactions seem largely unrelated to PCD *per se*, at least in our experimental conditions (e.g. amount of resources and duration of cycles). In fact these high-density dynamics are almost unchanged when PCD is shut off entirely (Figure 5). Overall, the interplay of these population dynamics at low and high density led to the maintenance of the PCD-inducing strain at high frequency. As a consequence, the mixed populations continued to exhibit marked decline following exposure to stress, even after 13 cycles of salinity rise.

In conclusion, we presented the first experimental evidence for evolutionary maintenance of programmed cell death in a unicellular organism, over many generations and multiple cycles of inductions by environmental stress. Our ability to distinguish genotypes in mixtures and track their population densities throughout the experiment allowed us to link selection and demography in a non-trivial ecological context. We could thus demonstrate that the decline-rebound population dynamics of PCD led to strong frequency fluctuations within a cycle, while competition at high density largely explained stable frequency across cycles and long-term coexistence.

## Notes

https://doi.org/10.5281/zenodo.12580685

### Competing Interest Statement

The authors have declared no competing interest.

### Summary of Updates

The introduction and discussion have been rewritten to clarify the aims of this study. Statistical analyses have been sumarised and the complete methods were moved to the electronic supplementary material.

## References

1. Crow JF, Kimura M. 1970 An Introduction to Population Genetics.

2. Darwin C. 1859 On the origin of species. John Murray, London.

3. De Jong G. 1994 The Fitness of Fitness Concepts and the Description of Natural Selection. Q. Rev. Biol. 69.

4. Lenormand, T., Rode, N., Chevin, L. M., & Rousset F. 2016 Valeur sélective: définitions, enjeux et mesures. In Biologie évolutive, pp. 655–675.

5. Chevin LM. 2011 On measuring selection in experimental evolution. Biol. Lett. 7, 210–213. (doi:10.1098/rsbl.2010.0580)

6. Lenski RE. 1991 Quantifying fitness and gene stability in microorganisms. In *Biotechnology (Reading*, Mass.), pp. 173–192. Butterworth-Heinemann. (doi:10.1016/b978-0-409-90199-3.50015-2)

7. Lenski RE, Rose MR, Simpson SC, Tadler SC. 1997 Long-term experimental evolution in. J. Evol. Biol. 10, 743. (doi:10.1007/s000360050052)

8. Fisher R. A., Ford E. B. 1947 The spread of a gene in natural conditions in a colony of the moth Panaxia dominula L. Heredity (Edinb). 1, 143:174. (doi:10.1021/ac60010a024)

9. Cain AJ, Sheppard PM. 1954 Natural Selection in Cepaea. Genetics 39, 89–116. (doi:10.1093/genetics/39.1.89)

10. Gallet R, Cooper TF, Elena SF, Lenormand T. 2012 Measuring selection coefficients below 10-3: Method, Questions, and Prospects. Genetics 190, 175–186. (doi:10.1534/genetics.111.133454)

11. Rescan M, Grulois D, Aboud EO, de Villemereuil P, Chevin L-M. 2021 Predicting population genetic change in an autocorrelated random environment: Insights from a large automated experiment. PLOS Genet. 17, e1009611. (doi:10.1371/journal.pgen.1009611)

12. Burke MK, Dunham JP, Shahrestani P, Thornton KR, Rose MR, Long AD. 2010 Genome-wide analysis of a long-term evolution experiment with Drosophila. Nature 467, 587–590. (doi:10.1038/nature09352)

13. Burny C, Nolte V, Dolezal M, Schlötterer C. 2022 Genome-wide selection signatures reveal widespread synergistic effects of two different stressors in Drosophila melanogaster. Proc. R. Soc. B Biol. Sci. 289. (doi:10.1098/rspb.2022.1857)

14. Levy SF, Blundell JR, Venkataram S, Petrov DA, Fisher DS, Sherlock G. 2015 Quantitative evolutionary dynamics using high-resolution lineage tracking. Nature 519, 181–186. (doi:10.1038/nature14279)

15. Gomulkiewicz R, Holt RD. 1995 When does evolution by natural selection prevent extinction? Evolution (N. Y). 49, 201–207.

16. Hendry AP. 2017 Eco-evolutionary Dynamics. Princeton University Press. (10.1515/9781400883080)

17. Ribeck N, Lenski RE. 2015 Modeling and quantifying frequency-dependent fitness in microbial populations with cross-feeding interactions. Evolution (N. Y). 69, 1313–1320. (doi:10.1111/evo.12645)

18. Estrela S, Libby E, Van Cleve J, Débarre F, Deforet M, Harcombe WR, Peña J, Brown SP, Hochberg ME. 2019 Environmentally Mediated Social Dilemmas. Trends Ecol. Evol. 34, 6–18. (doi:10.1016/j.tree.2018.10.004)

19. De Meester L, Vanoverbeke J, Kilsdonk LJ, Urban MC. 2016 Evolving Perspectives on Monopolization and Priority Effects. Trends Ecol. Evol. 31, 136–146. (doi:10.1016/j.tree.2015.12.009)

20. MacArthur RH. 1962 Some Generalized Theorems of Natural Selection. Proc. Natl. Acad. Sci. 48, 1893–1897. (doi:10.1073/pnas.48.11.1893)

21. Engen S, Lande R, Sæther BE. 2013 A quantitative genetic model of r- and K-selection in a fluctuating population. Am. Nat. 181, 725–736. (doi:10.1086/670257)

22. Sæther BE, Visser ME, Grøtan V, Engen S. 2016 Evidence for r-and K-selection in a wild bird population: A reciprocal link between ecology and evolution. Proc. R. Soc. B Biol. Sci. 283. (doi:10.1098/rspb.2015.2411)

23. Leung C, Grulois D, Chevin L-M. 2022 Plasticity across levels: relating epigenomic, transcriptomic, and phenotypic responses to osmotic stress in a halotolerant microalga. Mol. Ecol.

24. Zeballos N, Grulois D, Leung C, Chevin L-M. 2023 Acceptable loss: Fitness consequences of salinity-induced cell death in a halotolerant microalga. Am. Nat. 201. (10.1086/724417)

25. Ameisen JC. 2002 On the origin, evolution, and nature of programmed cell death: a timeline of four billion years. Cell Death Differ. 9, 367–393. (doi:10.1038/sj/cdd/4400950)

26. Bidle KD. 2016 Programmed Cell Death in Unicellular Phytoplankton. Curr. Biol. 26, R594–R607. (doi:10.1016/j.cub.2016.05.056)

27. Deponte M. 2008 Programmed cell death in protists. Biochim. Biophys. Acta 1783, 1396–1405. (doi:10.1016/j.bbamcr.2008.01.018)

28. Orellana M V, Pang WL, Durand PM, Whitehead K, Baliga NS. 2013 A Role for Programmed Cell Death in the Microbial Loop. PLoS One 8, e62595. (doi:10.1371/journal.pone.0062595)

29. Durand PM, Rashidi A, Michod RE. 2011 How an organism dies affects the fitness of its neighbors. Am. Nat. 177, 224–232. (doi:10.1086/657686)

30. Reece SE, Pollitt LC, Colegrave N, Gardner A. 2011 The meaning of death: Evolution and ecology of apoptosis in protozoan parasites. PLoS Pathog. 7, 1–9. (doi:10.1371/journal.ppat.1002320)

31. Rescan M, Grulois D, Ortega-Aboud E, Chevin LM. 2020 Phenotypic memory drives population growth and extinction risk in a noisy environment. *Nat*. Ecol. Evol. 4, 193–201. (doi:10.1038/s41559-019-1089-6)

32. Ben-Amotz A, Polle JEW, Subba Rao DV. 2009 The Alga Dunaliella Biodiversity, Physiology, Genomics and Biotechnology.

33. Slee EA, Adrain C, Martin SJ. 2001 Executioner Caspase-3, -6, and -7 Perform Distinct, Non-redundant Roles during the Demolition Phase of Apoptosis. J. Biol. Chem. 276, 7320–7326. (doi:10.1074/jbc.M008363200)

34. Kasuba KC, Vavilala SL, D’Souza JS. 2015 Apoptosis-like cell death in unicellular photosynthetic organisms - A review. Algal Res. 12, 126–133. (doi:10.1016/j.algal.2015.07.016)

35. Danial NN, Korsmeyer SJ. 2004 Cell Death: Critical Control Points. Cell 116, 205–219. (doi:10.1016/S0092-8674(04)00046-7)

36. Jacob, S. W., & Herschler R. 1986 Pharmacology of DMSO. Cryobiology 23(**1**), 14–27.

37. Venables WN, Ripley BD. 2002 Modern Applied Statistics with S. Springer, New York.

38. Van Buuren S, Groothuis-Oudshoorn K. 2011 mice: Multivariate imputation by chained equations in R. J. Stat. Softw. 45, 1–67. (doi:10.18637/jss.v045.i03)

39. Kot M. 2001 Elements of Mathematical Ecology. Cambridge University Press.

40. Violle C, Nemergut DR, Pu Z, Jiang L. 2011 Phylogenetic limiting similarity and competitive exclusion. Ecol. Lett. 14, 782–787. (doi:10.1111/j.1461-0248.2011.01644.x)

41. Gause GF. 1934 Experimental analysis of Vito Volterra’s mathematical theory of the struggle for existence. Science (80-. ). 79, 16–17. (doi:10.1126/science.79.2036.16-a)

42. MacArthur R, Levins R. 1967 The limiting similarity, convergence, and divergence of coexisting species. Am. Nat. 101, 377–385.

43. Ginzburg M, Ginzburg BZ. 1985 Ion and glycerol concentrations in 12 isolates of Dunaliella. J. Exp. Bot. 36, 1064–1074. (doi:10.1093/jxb/36.7.1064)

