## Supplementary material for "Linking selection to demography in experimental evolution of active death in a unicellular organism"

Running title: Selection on population decline

Nathalie Zeballos[1], Océane Rieu[1], Stanislas Fereol[1], Christelle Leung[1], and Luis-Miguel Chevin[1]

[1] CEFE, Univ Montpellier, CNRS, EPHE, IRD, Montpellier, France

Keywords: population decline, natural selection, competition, experimental evolution, environmental stress, programmed cell death.

This is the supplementary material regarding this article.

.

### Supplementary Material

#### 1. DNA confirmation

At the end of the 13 cycles of the assay, we genetically checked for the presence of both strains within each evolved population. Specifically, we amplified one mitochondrial (333 pb fragment using the primers DsMt1-For [5'-GGTTAGTCATAGTTGGAGGT-3'] and DsMt1-Rev [5'-GAAAACCTAACATGGCTAAGC-3']) and one chloroplastic (372 bp fragment using the primers DsChl1-For [5'-TTTAGGCGAATCCATAAGAG-3'] and DsChl1-Rev [5'-CCAAGCAGGTGAATTAGCTTTG-3']) locus from [1], specific to strains A and C, respectively. DNA was extracted from *c.* $1.10^6$ cells using Nucleospin plant II (Macherey-Nagel) and amplified as followed: PCR was conducted in a total volume of 20 µL, including 10 µL of MasterMix Phusion with HF Buffer 2X (ThermoScientific), 2 µL of DNA, 10 µM of forward and reverse primers and 5 µL of pure $H_2O$. Amplification cycle consisted in: 30 s initial denaturation at 92˚C; 45 cycles of 15 s at 92°C, 15 s at 54°C, 30 s at 68°C; and a final extension at 68°C for 5 min. Presence of each strain has been confirmed in all long-term mixtures by amplicon migration on agarose gels (Fig. S3).

#### 2. Long-term competition over successive hyper-osmotic shocks

To investigate the outcome of competition between the declining and non-declining strain over a longer time span, we maintained long-terms mixtures in their cycling treatments (4-10d or 7-7d in intermediate-high salinities), over a total of 13 cycles. From cycle 6 to 11 included, populations densities were measured only before and after dilution, and on the day following each hyper-osmotic shock. For the two last cycles, we followed the usual measurement pattern, where demography was tracked daily for 7 or 10 days following the hyper-osmotic shock. Since we no longer had the corresponding monocultures needed for our frequency estimation method, we instead used as reference the mean values of the monocultures at the same day of cycle 5, i.e the last cycle with monocultures flasks (Fig. S5).

#### 3. Testing DMSO effect in PCD inhibition

We performed additional test to verify that DMSO, used as solvent for our PCD inhibitor, was not responsible for the reduction in PCD observed in our experiments shown in Fig. 5. We first acclimated 10 replicates monocultures of strain A for 4 days at 2.4M NaCl, with an initial density set at $N_0 = 50{,}000$ cells/ml, in total culture volume of 10ml. We then applied three conditions to 500,000 cells per flask before a hyper-osmotic stress experiment: 1) a standard treatment corresponding to the low salinity condition in our main experiment; 2) a DMSO treatment where 10 µl of pure DMSO were added, without inhibitor; 3) and a DMSO + inhibitor treatment, where cells were treated with 10 µM Z-VAD(Ome)-FMK (CellSignaling) caspase inhibitor diluted in DMSO, corresponding to 10 µl of the solution. All flasks were then incubated in the dark for 30min at 24°C, before being subjected to a hyper-osmotic shock at 4M NaCl. Population densities were measured 1 and 4 hours after the hyper-osmotic transfer, and then daily tracked over 4 days. To estimate the intensity of the decline, we applied a GLM to population density 1 hour post-transfer, with cytometer cell concentration at day 0 as response variable, a log link function, negative binomial error structure, and log expected initial population density as offset (following [1–3]). Results are shown in Table S6. This assay confirmed that the inhibition of the decline is due to the caspase activity inhibitor, and not to the solvent DMSO. Actually, DMSO without inhibitor triggered higher population reduction than the hyper-osmotic shock alone, indicating that the effect of the inhibitor described in the main text is conservative.

#### 4. Inferring genotype frequencies and densities in mixtures from phenotypic measurements

##### Relative frequencies of strains

Our focal strains A and C differed in their natural yellow and red fluorescence in most mixed populations, but with distributions that overlapped to some extent (Fig. S1), so we could not use hand-designed cytometry gates to directly count the numbers of each strain. Instead, we estimated strain frequencies in mixtures using a custom-made maximum likelihood method that applies to any mixture of genotypes with different phenotypic distributions for continuous traits (we compared the two methods, see Material S6 and Fig. S2). We assumed that the pair of traits (yellow and red fluorescence) followed a bivariate Gaussian distribution in each monoculture, with mean vector $\mu$ and covariance matrix $\Sigma$ specific to each strain. The distribution of these traits in each mixed population was then treated as a mixture of Gaussians from the corresponding monocultures (sampled on the same day), with frequency $p$ for the declining strain A and *1-p* for the non-declining strain C. As all points were measured independently within each assay, we summed their log-likelihoods, and further transformed frequencies to the logit scale $u = \ln(\frac{p}{1-p})$ to facilitate maximization, leading to

$$lnLik = \sum \log\left(\frac{1}{1+e^{-u}} \times f_A(\mu_A, \Sigma_A) + \frac{e^{-u}}{1+e^{-u}} \times f_C(\mu_C, \Sigma_C)\right). \quad \text{(Eq. S1)}$$

We obtained the estimator of $\hat{u}_i$ by maximizing log-likelihood for each isogenic line $i$. We also extracted the curvature $c_i$ (Hessian matrix) of the log-likelihood function at its maximum to estimate uncertainty as $V(\hat{u}_i) = -1/c_i$, relying the asymptotic normal approximation of likelihood [4,5]. We then computed the 95% confidence interval endpoints on the logit scale for each estimator $\hat{u}_i$ as

$$CI_{95\%}(u_i) = \left[\hat{u}_i - 1.96\sqrt{V(\hat{u}_i)}; \hat{u}_i + 1.96\sqrt{V(\hat{u}_i)}\right]. \quad \text{(Eq. S2)}$$

Isogenic lines of strains A and C differed in how well they could be distinguished in mixtures, so the mean frequency $p$ was estimated with variable uncertainties depending on the mixture. We corrected it by placing more weight on more precise estimates with the inverse-variance weighting method, which is commonly used in meta-analyses (when sample sizes differ across studies) [6], and whenever there are known sources variation in uncertainty across estimates

[7]. Accordingly, the mean $\hat{u}$ over the 10 mixtures of isogenic lines $i$ was obtained for each treatment and measurement as

$$\hat{u}_{mean} = \frac{\sum_i c_i \times \hat{u}_i(p)}{\sum_i c_i} \quad \text{(Eq. S3)}$$

and the error variance of $\hat{u}_{mean}$ as

$$V(\hat{u}_{mean}) = -\frac{1}{\sum_i c_i}. \quad \text{(Eq. S4)}$$

We then computed the 95% confidence interval endpoints on the logit scale for $\hat{u}_{mean}$ by inserting eqs. (B3) and (B4) into the eq. (B2), leading to

$$CI_{95\%}(u_{mean}) = \left[\hat{u}_{mean} - 1.96\sqrt{V(\hat{u}_{mean})}; \hat{u}_{mean} + 1.96\sqrt{V(\hat{u}_{mean})}\right]. \quad \text{(Eq. S5)}$$

For further analyses, we back-transformed the frequency of A in each replicate to the arithmetic scale using the delta method for bias correction [8] as

$$\hat{p}_i = \frac{1}{1+e^{-\hat{u}_i}} + \frac{V(\hat{u}_i)}{2} \times g''(\hat{u}_i) \quad \text{(Eq. S6)}$$

where

$$g''(\hat{u}_i) = \frac{-e^{\hat{u}_i}\,(e^{\hat{u}_i}-1)}{(1+e^{\hat{u}_i})^3}.$$

is the second derivative of the logistic function (inverse of the logit function). We back-transformed the confidence interval from eq. (B2) by applying the logistic function to its end points. For the mean frequency over the 10 isogenic lines (in each condition and day, Fig. 2), we also used the delta method by combining eqs. (B3) and (B4) with eq. (B6), leading to

$$\hat{p}_{mean} = \frac{1}{1+e^{-\hat{u}_{mean}}} + \frac{V(\hat{u}_{mean})}{2} \times g''(\hat{u}_{mean}). \quad \text{(Eq. S7)}$$

The confidence interval for the mean frequency was then obtained by applying the logistic function to its end points obtained with the eq. (B5). Finally, the error variance for the mean frequency $\hat{p}_{mean}$ was obtained with the delta method [8] as

$$V(\hat{p}_{mean}) = \frac{e^{2\hat{u}_{mean}}}{(1+e^{\hat{u}_{mean}})^4} \times V(\hat{u}_{mean}). \quad \text{(Eq. S8)}$$

#### Inferred strain densities over the first 5 cycles

From these estimates of relative frequencies, we inferred the densities of each strain in the mixtures as $\hat{p}_i \times N_{tot}$, with $\hat{p}_i$ the estimated frequency of each strain in replicates $i$ and $N_{tot}$ the total population density (in cells/ml) measured in the flask. We then computed the daily mean inferred density of strain A in the long-term mixtures over the 5 cycles as

$$N_{A,cyc} = E_{cyc}\left(N_{tot,mean}\hat{p}_{mean}\right) = E_{cyc}\left(N_{tot,mean}\right) \times E_{cyc}(\hat{p}_{mean}), \quad \text{(Eq. S9)}$$

where $E_{cyc}$ denotes an expectation over cycles, $N_{tot,mean}$ and $\hat{p}_{mean}$ are respectively the total population density and frequency of strain A averaged over the 10 isogenic lines within a cycle, and the second member assumes that $N_{tot,mean}$ and $\hat{p}_{mean}$ are independent across cycles. The density of strain C was obtained similarly as $N_{C,cyc} = E_{cyc}\left(N_{tot,mean}\right) \times (1 - E_{cyc}(\hat{p}_{mean}))$. Accordingly, the error variance of $N_{tot,mean}\hat{p}_{mean}$ over the 10 isogenic lines per condition within a cycle was

$$V\left(N_{tot,mean}\hat{p}_{mean}\right) = V\left(N_{tot,mean}\right) \times V(\hat{p}_{mean}) + V\left(N_{tot,mean}\right) \times E(\hat{p}_{mean})^2 + V(\hat{p}_{mean}) \times E\left(N_{tot,mean}\right)^2 \quad \text{(Eq. S10)}$$

(product of two independent random variables). We then computed the 95% confidence interval of the inferred densities over the 5 cycles using the law of the total variance, $V_{cyc}\left(N_{A,cyc}\right) = E_{cyc}\left(V\left(N_{tot,mean}\hat{p}_{mean}\right)\right) + \frac{V_{cyc}(E(N_{tot}\hat{p}_{mean}))}{nb\ cycles}$, where $E_{cyc}$ and $V_{cyc}$ denote expectations and variances across cycles, and similarly for strain C.

#### 5. Analyses of density-dependent competition

We estimated the strength of density dependent competition through the relationship between *per-capita* growth rate (over a short time interval) and population density in mixtures. However, population densities were estimated with error from small cytometer samples [9]. Estimating the density of each strain in a mixture further required estimating the relative frequency of each strain using our mixture distribution approach (Material S4, eqs. S1-S8). This approach not only involves uncertainty in the estimator $\hat{u}_i$ of logit frequency of strain A, but this uncertainty also differs among mixtures of different clones of strains A and C. To account for all these sources of uncertainty, we simulated 1000 datasets, where the frequency $u_{i,simulated}$ of strain A (on the logit scale) in mixture *i* was randomly sampled from a gaussian distribution with mean $\hat{u}_i$ (the estimator of $u_i$), and standard deviation $\sqrt{V(u_i)}$. We then back-transformed the strain frequencies to the arithmetic scale using the inverse logit function. We also simulated the total number of cells in mixed populations by sampling from a negative binomial distribution with mean $\widehat{N}_\iota$ and dispersion parameter = 10 (consistent with estimations from GLMs, e.g. [9]). We then computed the simulated density of each strain in mixed cultures for each simulated dataset, as previously explained. From this, the *per-capita* growth rate per day of strain A in mixed populations (for each simulated dataset) was computed as

$$r_A = \frac{\widehat{N}_{A,t} - \widehat{N}_{A,t-\tau}}{\tau \, \widehat{N}_{A,t-\tau}} \qquad \text{(Eq. S11)}$$

with $\tau$ the time interval between successive measurements (mostly 1 day, sometimes 2 or 3 days), and similarly for strain C. We restricted the analysis to times $t$ after day 4 (included) and population densities $\widehat{N}_t >$ 5.10^4 cells/ml for both strains, so as to exclude the initial phase of decline-rebound where the relationships between population growth and size are not driven by density dependence. We also focused only on clone mixtures that allowed for clear distinction of both strains, such that the uncertainty in frequency estimation was low ($CI_{95\%}(u_i)$ width $<$ 0.5 on the logit scale).

Finally, we fitted linear models (LM) with the *per-capita* growth rate as response variable, and the densities $\widehat{N}_{t-\tau}$ of both strains or the total densities in mixtures as fixed effects (see Table S1 for details), on each of the 1000 simulated dataset. We computed overall estimates with standard error and p-values over the 1000 LMs using Rubin's rules (assuming normal distributions) [10]. We also computed predictions over the 1000 LMs and combined them to produce an overall fitted line [11] (Fig. 4). These linear models correspond to the classic Lotka-Volterra model of linear density-dependent competition, so we could check the coexistence condition [12]. However, the relationship between the *per-capita* growth rate and population density need not be linear, so we also fitted semi-parametric models (cubic splines) using generalized additive models (GAM) (see Table 1 for formula).

#### 6. Calibrating and testing the efficiency of the frequency estimation method

We tested the efficiency of our estimation method using a calibration assay. We took 8 of the 10 monocultures per strain at high salinity (cycle 3), and prepared the corresponding mixed populations over a range of expected frequencies (from 5% to 95%). These proportions were based on volumes for practical reasons, so the expected frequencies were deduced from the densities of the monocultures used to prepare the mixtures. We then counted cell densities in these freshly prepared flasks using our flow cytometer. The proportions of cells from strain A and strain C estimated using restrictive hand-drawn gates (see Figure S2A) were compared to the those obtained through our estimation method (Figure S2B). We then simulated mixed populations by virtually sampling 10,000 cells from the datasets of strain A and strain C monocultures, at different frequencies (note that for a flask at equilibrium stationary phase, less than 20,000 events are recorded in the 30s of flow cytometry). We then applied our estimation method on these simulated mixtures with predefined frequencies (Figure S2C).

#### Supplementary Tables

**Table S1. Model selection for density dependence, based on linear models (LMs) or generalised additive models' (GAMs).**

| **LMs** | **Variable response: $r_{strain\,1}$**<br>**Fixed effects:** | **AIC**<br>$r_{strain\,A}$ | AIC<br>$r_{strain\,C}$ |
|---|---|---|---|
| **Model 1** | $N_{strain\,1,t-\tau} + N_{strain\,2,t-\tau}$ | **432.61** | **442.63** |
| **Model 2** | $N_{Total,t-\tau}$ | 444.18 | 447.77 |
| **GAMs** | **Variable response: $r_{strain\,1}$**<br>**Fixed effects:** | AIC $r_{strain\,A}$ | AIC<br>$r_{strain\,C}$ |
| **Model 1** | $s(N_{strain\,1,t-\tau}) + s(N_{strain\,2,t-\tau})$ | **418.85** | **433.06** |
| **Model 2** | $N_{Total,t-\tau}$ | 438.79 | 445.99 |

In the GAMs formula, $s$ is a function used in definition of smooth terms. Akaike information criterion was averaged over 1000 LMs (estimated on resimulated datasets to account for uncertainty). The best model based on AIC is shown in bold.

**Table S2. Population decline rate against number of salinity cycles**.

| | Estimate | Std. Error | z value | Pr(>\|z\|) |
|---|---|---|---|---|
| Intercept (A+C control mixtures in cycle 1 in 4-10d fluctuations) | -0.434 | 0.076 | -5.683 | 1e-08 *** |
| Cycle | -0.008 | 0.023 | -0.337 | 0.736 |
| Strain_Amonocultures | -0.866 | 0.108 | -8.026 | 1e-15 *** |
| Strain_long-termA+Cmixtures | -0.158 | 0.087 | -1.802 | 0.0715 . |
| Strain_Cmonocultures | 0.496 | 0.108 | 4.601 | 4e-6 *** |
| Fluct_7-7 | 0.102 | 0.108 | 0.949 | 0.343 |
| Cycle:Strain_Amonocultures | -0.065 | 0.033 | -2.010 | 0.044 * |
| Cycle:Strain_long-termA+Cmixtures | 0.003 | 0.024 | 0.110 | 0.913 |
| Cycle:Strain_Cmonocultures | -0.004 | 0.033 | -0.125 | 0.901 |
| Cycle:fluct_7-7 | 0.000 | 0.033 | -0.010 | 0.992 |
| Strain_Amonocultures:fluct_7-7 | 0.270 | 0.149 | 1.818 | 0.069 . |
| Strain_long-termA+Cmixtures:fluct_7-7 | -0.118 | 0.124 | -0.951 | 0.341 |
| Strain_Cmonocultures:fluct_7-7 | -0.089 | 0.153 | -0.585 | 0.558 |
| Cycle:Strain_Amonocultures:fluct_7-7 | 0.005 | 0.043 | 0.108 | 0.914 |
| Cycle:Strain_long-termA+Cmixtures:fluct_7-7 | 0.008 | 0.033 | 0.227 | 0.821 |
| Cycle:Strain_Cmonocultures:fluct_7-7 | 0.005 | 0.046 | 0.117 | 0.907 |

Generalised linear model using density measured 1 hour after the salinity rise as response variable, and the logarithm of expected initial population density as offset. We tested the effect of strain cultures identity (monocultures or mixtures), the cycle duration as factor, and the number of cycles as a continuous variable. The interaction of other factors with the number of cycles represents the effects of these factors over the successive salinity rises. p-value: ***: <0.001 ; **: <0.01 ; : <0.05 ; **.**: <0.1.

**Table S3. Effect of self-density and competitor density on *per-capita* growth rate for both strains A and C (LM 1).**

| $r_A$ | Estimate | Std. Error | Statistic | Df | Pr(>\|t\|) |
|---|---|---|---|---|---|
| Intercept | 1.29 | 1.58E-01 | 8.17 | 102.62 | 8.35E-13 *** |
| **$N_{A, t-\tau}$** | -3.10E-06 | 5.29E-07 | -5.86 | 105.81 | 5.34E-08 *** |
| **$N_{C, t-\tau}$** | -7.71E-07 | 6.03E-07 | -1.28 | 114.95 | 0.20 * |
| **$r_C$** | **Estimate** | **Std. Error** | **Statistic** | **Df** | **Pr(>\|t\|)** |
| Intercept | 1.15 | 1.53E-01 | 7.50 | 112.49 | 1.55E-11 *** |
| **$N_{C, t-\tau}$** | -2.85E-06 | 5.78E-07 | -4.93 | 128.37 | 2.48E-06 *** |
| **$N_{A, t-\tau}$** | -1.14E-06 | 4.79E-07 | -2.39 | 134.43 | 1.83E-02 *** |

We applied a linear model (LM 1) on the *per-capita* growth rate as response variable, where fixed effects were self-density and the density of the competitor. Growth rates were computed on 1000 simulated datasets based on the estimated strain densities within mixtures. We calculated the mean for the estimates and their variance over the 1000 LM according to Rubin's rules. p-value: ***: <0.001 ; **: <0.01 ; : <0.05 ; .: <0.1.

**Table S4. Effect of total density in the mixture on *per-capita* growth rate (LM 2).**

| $r_A$ | Estimate | Std. Error | Statistic | Df | Pr(>\|t\|) |
|---|---|---|---|---|---|
| Intercept | 1.28 | 1.62E-01 | 7.92 | 104.03 | 2.71E-12 *** |
| $\mathbf{N_{total,\ t-\tau}}$ | -2.09E-06 | 3.58E-07 | -5.82 | 110.58 | 5.77E-08 *** |
| $r_C$ | **Estimate** | **Std. Error** | **Statistic** | **Df** | **Pr(>\|t\|)** |
| Intercept | 1.16 | 1.55E-01 | 7.45 | 113.16 | 2.01E-11 *** |
| $\mathbf{N_{total,\ t-\tau}}$ | -1.88E-06 | 3.41E-07 | -5.49 | 121.81 | 2.18E-07 *** |

We applied a linear model (LM2) on the *per-capita* growth rate as response variable and total density in the mixture as fixed effects. Growth rates were computed based on measured densities in monocultures and estimated strain densities within mixtures. p-value: ***: <0.001 ; **: <0.01 ; : <0.05 ; **.**: <0.1.

**Table S5. Effect of initial density and PCD inhibitor treatment on strain A monocultures rebound.**

| | Estimate | Std. Error | z value | Pr(>\|z\|) |
|---|---|---|---|---|
| Intercept (Strain A monocultures without PCD inhibitor starting at N0= 25,000 cells/ml) | 7.620 | 0.125 | 61.064 | <2e-16 *** |
| Time | 0.746 | 0.034 | 21.976 | <2e-16 *** |
| Inhibitor_10 | 2.087 | 0.176 | 11.830 | <2e-16 *** |
| Initialdensity_50,000 | 0.871 | 0.153 | 5.699 | 1.20e-8 *** |
| Time:Inhibitor_10 | -0.318 | 0.048 | -6.622 | 3.55e-11 *** |
| Time:Initialdensity_50,000 | -0.031 | 0.042 | -0.754 | 0.451 |
| Inhibitor_10:Initialdensity_50,000 | 0.001 | 0.216 | 0.003 | 0.997 |
| Time:Inhibitor_10:Initialdensity_50,000 | -0.060 | 0.059 | -1.026 | 0.305 |

GLM on population densities on days 2 to 5 of strain A monocultures from the PCD inhibitor assay, corresponding to growth in the exponential phase (visually detected as in [3]). Our main interest is the effect of Time and its interactions with other factors, which represent effects of these factors (here PCD inhibitor doses and initial density) on maximal population growth rate. p-value: ***: <0.001 ; **: <0.01 ; : <0.05 ; .: <0.1.

**Table S6: Effect of DMSO on initial decline following hyper-osmotic shock (1-hour after hyper-osmotic transfer).**

| **Table A** | Estimate | Std. Error | z value | Pr (>\|z\|) | Population reduction (%) |
|---|---|---|---|---|---|
| Intercept (Standard hyper-osmotic shock) | -0.655 | 0.034 | -19.557 | 3.56e-85 *** | 66 |
| Treatment_DMSO | -0.274 | 0.047 | -5.784 | 7.30e-09 *** | 93 |
| Treatment_Inhibitor | 0.721 | 0.047 | 15.218 | 2.69e-52 *** | -7 |

GLM on strain A population density at day 0 relative to the expected initial density, from the DMSO assay. We tested the effect of DMSO on demographic decline on the logarithmic scale (from the log link function in the GLM, using log expected initial density as offset), and also reported the proportional reduction in population size (where negative values denote proportional increase). p-value: ***: <0.001 ;  **: <0.01 ; : <0.05 ;  **.**: <0.1.

#### Supplementary Figures

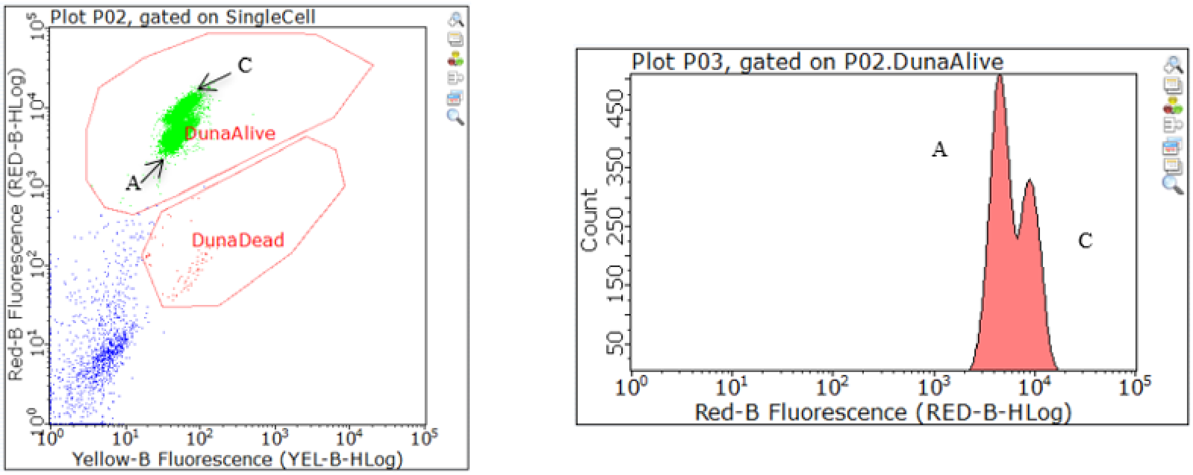


**Figure S1. Example of cytogram with raw data counts of a mixed strains A and C population.** (A) Dots from the two strain overlap but are still noticeable. (B) Gaussian distribution of dots included in the DunaAlive gate for red fluorescence trait in a mixture.

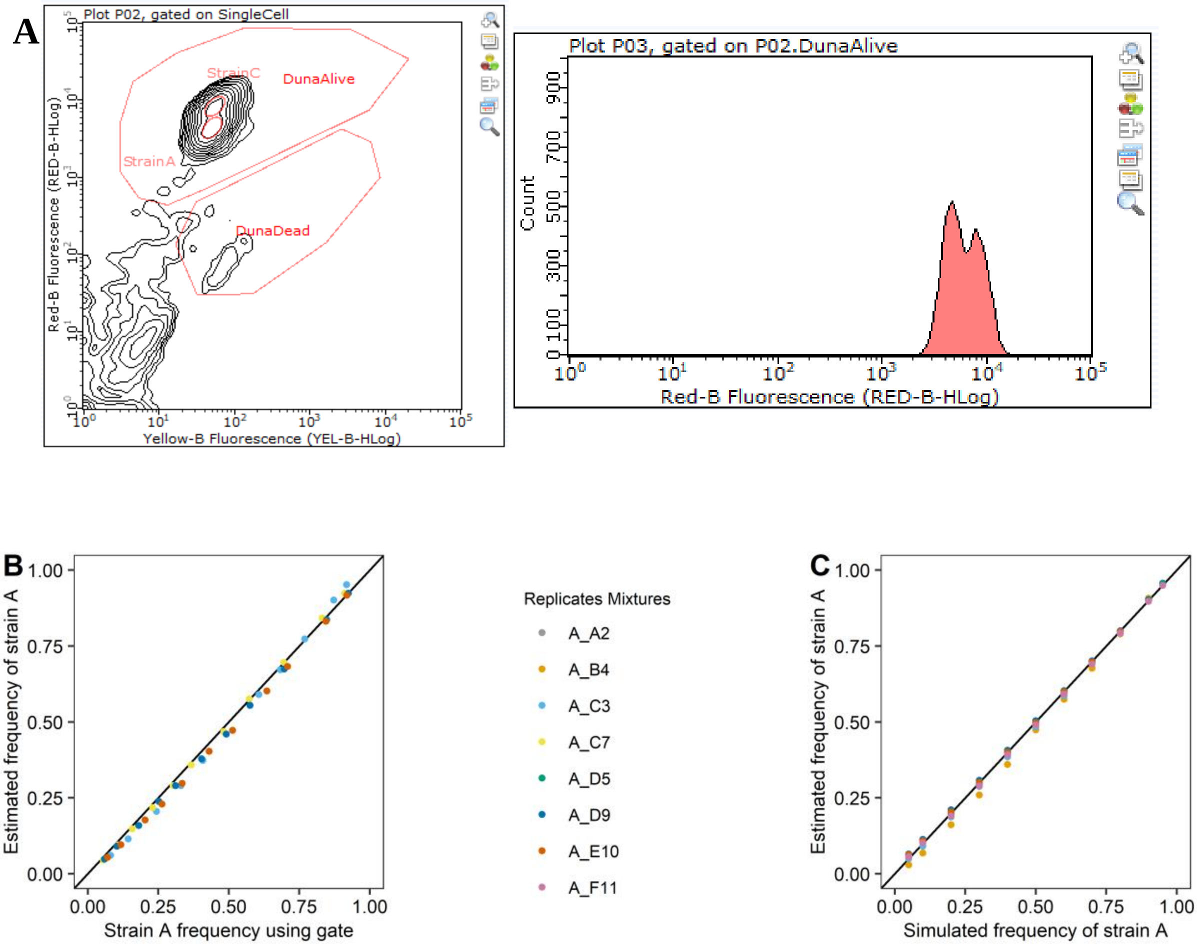


**Figure S2. Calibration assay for our estimation method of frequency in mixed populations.** (A) Restrictive handmade gates (small red ellipses) to discriminate strain A and C cells on the cytogram. This example shows a flask composed of populations that can be distinguished by red fluorescence. (B) Comparison between the frequency of strain A obtained by our estimation method, versus restrictive hand-drawn gates on the cytogram (4 replicates mixtures of well-distinguishable clones of strains A and C). (C) Same as B, but using simulated mixed populations (8 replicates mixtures).

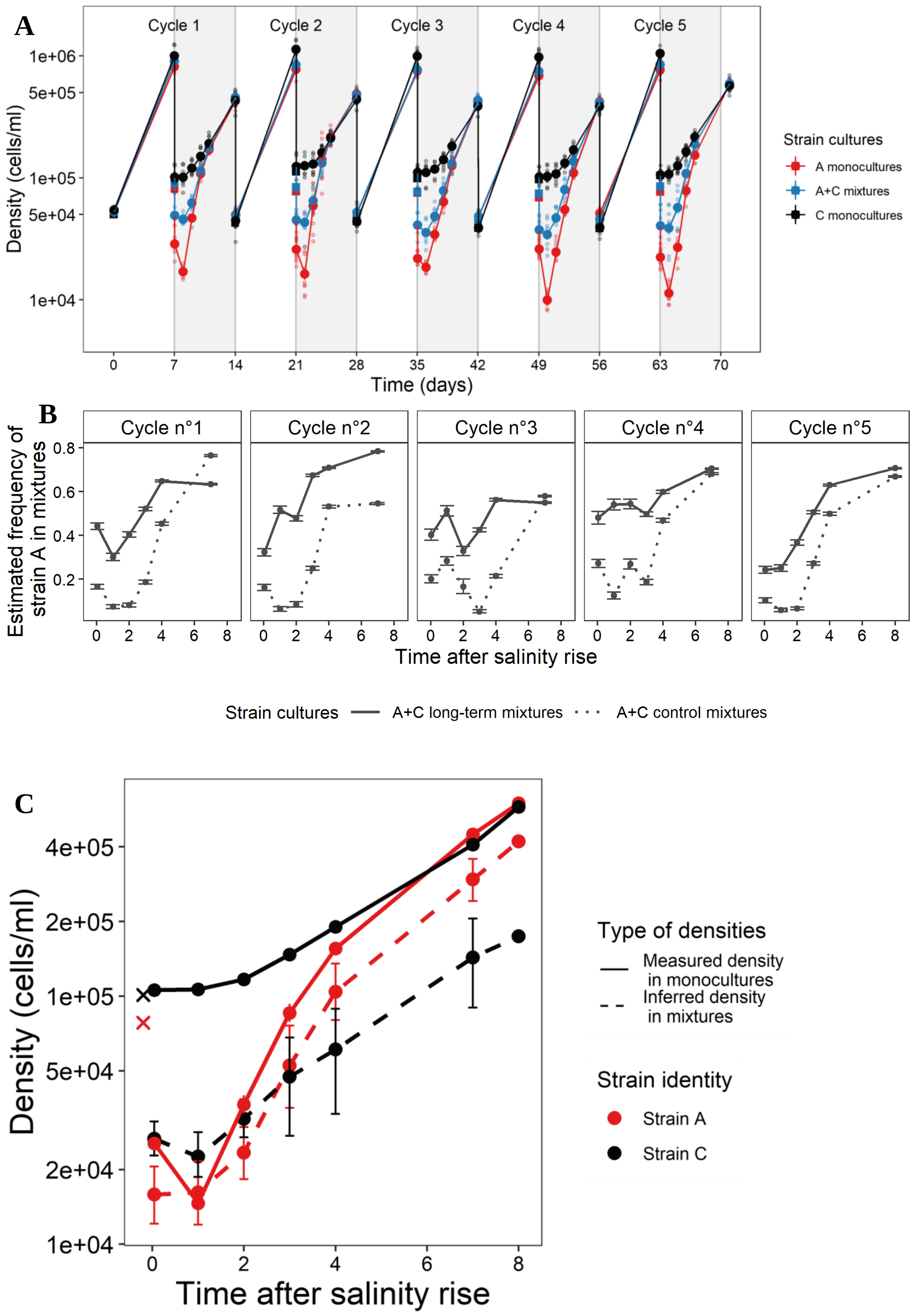


**Figure S3. 7-7d fluctuation cycle results**. (A) Demographic responses to successive salinity rises under 7-7d cycle. Population densities over the first five salinity cycles are shown for monocultures of strain A (red) and strain C (black), and long-term mixtures of A and C (blue). Small dots represent biological replicates (specific isogenic lines of each strain, or mixtures thereof), while large dots represent means over replicates (with SE not visible). Grey backgrounds correspond to the high salinity step in a given cycle. (B) Frequency of declining strain over 5 successive salinity rises. The proportion of mixed populations composed of strain A, as estimated from the cytometric traits of cells, is shown against time for long-term mixtures (solid lines), and control mixtures (dotted lines) that were freshly composed by mixing 50-50% of each strain before each salinity rise. Means and 95% confidence interval were obtained on the logit scale by inverse-variance weighting over the 10 population lines, to account for the variable uncertainty of estimates among biological replicates (isogenic lines of each strain and back-transformed to the arithmetic scale using the delta method. (C) Growth rates in monocultures versus mixtures. Mean population densities across replicates in monocultures (solid lines; direct measurements), and long-term mixtures (dashed lines; inferred from the estimate of relative frequencies), averaged over the first 5 high salinity steps, are shown for the 7-7d fluctuation cycle. For the monocultures, we computed the mean and standard error (not visible) over 10 isogenic lines. For the mixtures, we used inverse-variance-weighted mean and 95% confidence interval (computed with the delta method), to account for variable precision in estimation of strain A frequency among biological replicates. The crosses at time 0 correspond to the expected initial densities for monocultures, based on the known density before the 10% dilution. The dot for day 8 is because we measured the final day before transfer on day 8 instead of 7 for cycle 5.

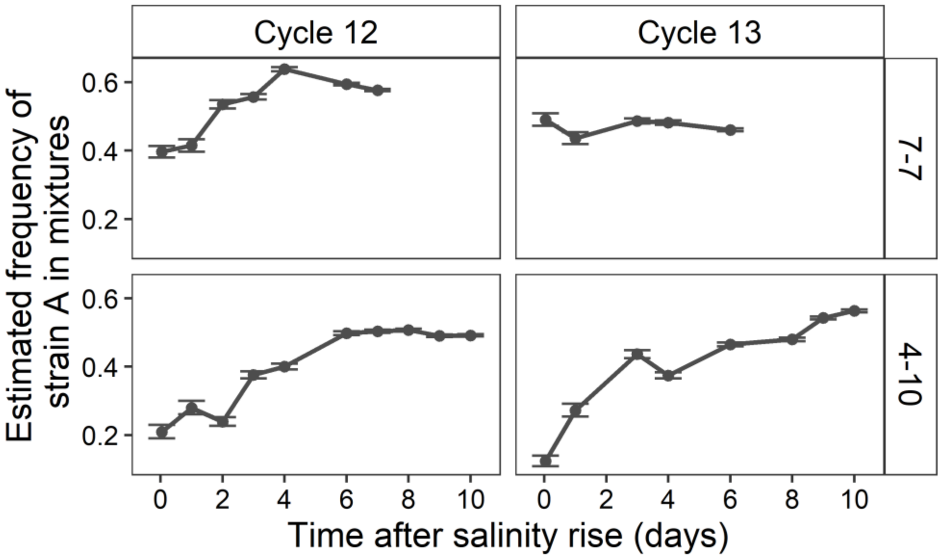


**Figure S4. Frequency of declining strain in cycles 12 and 13 in long-term mixtures.** The proportion of strain A in mixed populations, as estimated from the cytometric traits of cells, is shown against time for the last two cycles long-term mixtures, using the monocultures from cycle 5 as reference. Means and 95% confidence interval were obtained on the logit scale by inverse-variance weighting over the 10 population lines, to account for the variable uncertainty of estimates among biological replicates (isogenic lines of each strain), and back-transformed to the arithmetic scale using the delta method.

**A**

| Marker | Replicates mixtures + DsMt_1 (4-10d) | | | | | | | | | | Control + | Control + | Control - | X | Marker |
|---|---|---|---|---|---|---|---|---|---|---|---|---|---|---|---|
| Marker 100pb | A_A2 + C_A2 | A_B4 + C_B4 | A_C3 + C_C3 | A_C7 + C_C7 | A_D5 + C_D5 | A_D9 + C_D9 | A_E10 + C_E10 | A_F11 + C_F11 | A_G3 + C_G3 | A_H9 + C_H9 | A_A2 (rep 1) + DsMt_1 | A_A2 (rep 2) + DsMt_1 | C_A2 (rep 1) + DsChl1 | | Marker 100pb |

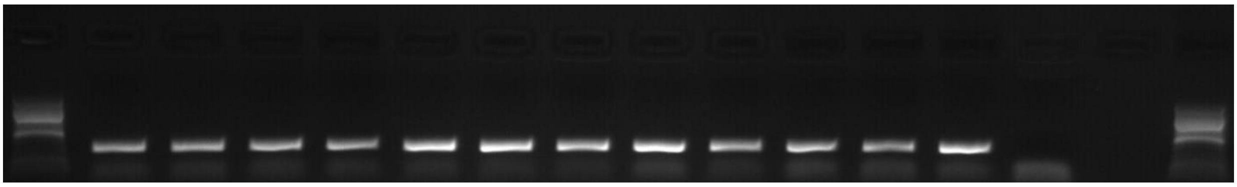

**B**

| Marker | Replicates mixtures + DsChl_1 (4-10d) | | | | | | | | | | Control + | Control + | Control - | Control + | Marker |
|---|---|---|---|---|---|---|---|---|---|---|---|---|---|---|---|
| Marker 100pb | A_A2 +C_A2 | A_B4 + C_B4 | A_C3 +C_C3 | A_C7 +C_C7 | A_D5 +C_D5 | A_D9 +C_D9 | A_E10 +C_E10 | A_F11 +C_F11 | A_G3 +C_G3 | A_H9 +C_H9 | C_A2 (4 months) + DsChl_1 | C_H9 (4 months) + DsChl1 | A_A2 (1 month) + DsMt_1 | C_A2 (1 month) + DsChl1 | Marker 100pb |

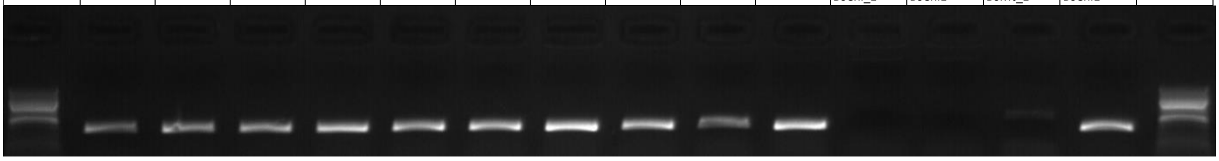

**C**

| Marker | Replicates mixtures + DsMt_1 (7-7d) | | | | | | | | | | Control + | Control - | Control - | Marker |
|---|---|---|---|---|---|---|---|---|---|---|---|---|---|---|
| Marker 100pb | A_A2 + C_A2 | A_B4 + C_B4 | A_C3 + C_C3 | A_C7 + C_C7 | A_D5 + C_D5 | A_D9 + C_D9 | A_E10 + C_E10 | A_F11 + C_F11 | A_G3 + C_G3 | A_H9 + C_H9 | A_B8 + DsMt_1 | C_B8 + DsChl1 | H2O + DsMt_1 | Marker 100pb |

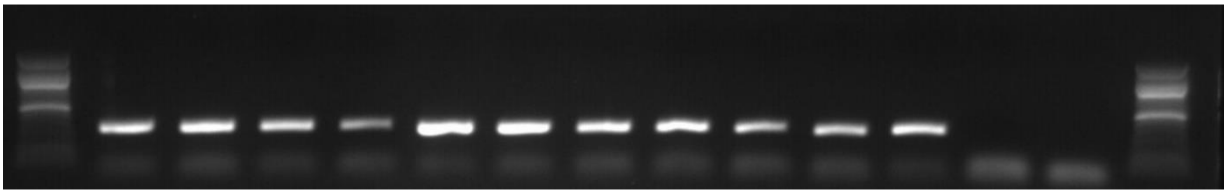

**D**

| Marker | Replicates mixtures + DsChl_1 (7-7d) | | | | | | | | | | Control + | Control - | Control - | Marker |
|---|---|---|---|---|---|---|---|---|---|---|---|---|---|---|
| Marker 100pb | A_A2 +C_A2 | A_B4 + C_B4 | A_C3 +C_C3 | A_C7 +C_C7 | A_D5 +C_D5 | A_D9 +C_D9 | A_E10 +C_E10 | A_F11 +C_F11 | A_G3 +C_G3 | A_H9 +C_H9 | C_B8 + DsChl_1 | A_B8 + DsMt_1 | H2O + DsChl_1 | Marker 100pb |

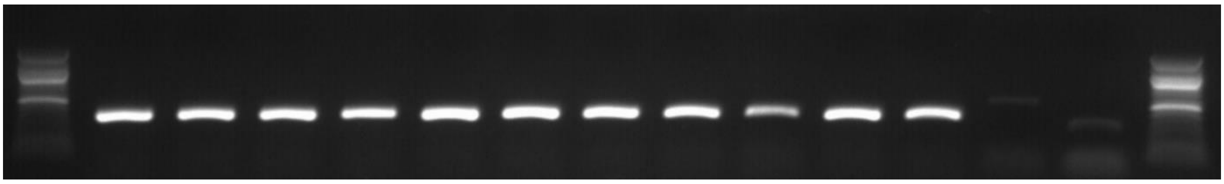

**Figure S5. Genetic confirmation of the presence of both strains in long-term mixtures after 13 cycles (26 weeks).** Amplicon migration on agarose gels of DsMt1 and DsChl1 locus, specific to strain A and C respectively. (A) Strain A presence (primer Ds_Mt1) on flasks under 4-10d cycle. (B) Strain C presence (primer Ds_Chl1) on flasks under 4-10d cycle. (C) Strain A presence (primer Ds_Mt1) on flasks under 7-7d cycle. (D) Strain C presence (primer Ds_Chl1) on flasks under 7-7d cycle. Positive and negative controls were also included (samples at the right position).

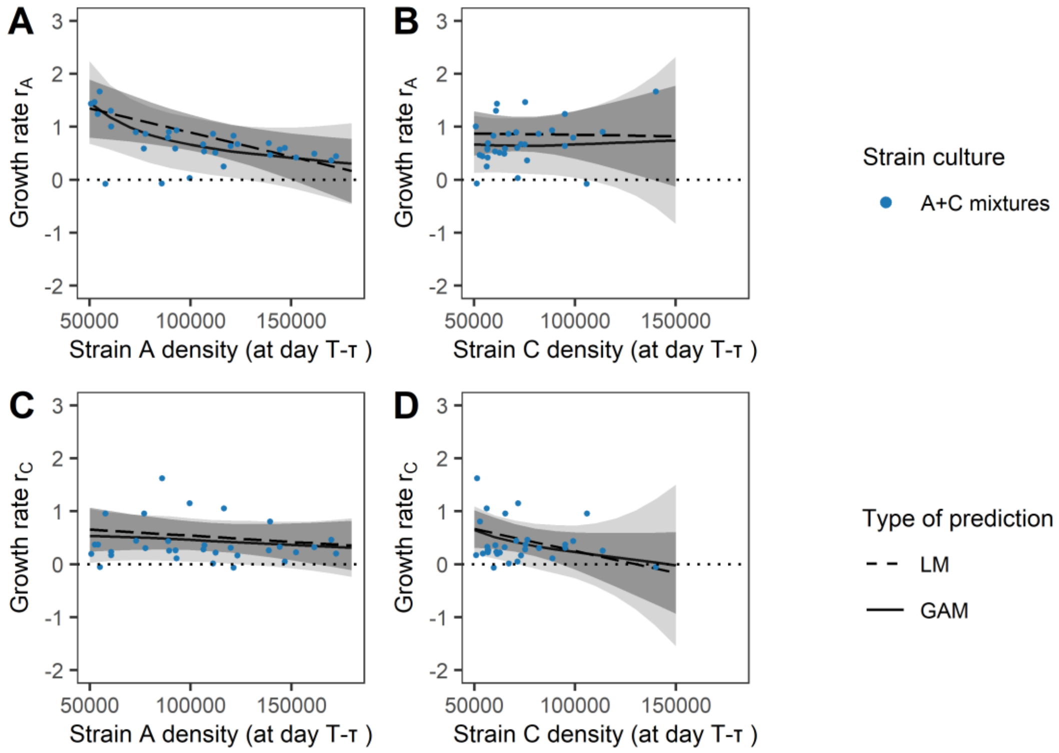


**Figure S6. Intra- and inter-strain competition and population growth in 7-7d fluctuation cycles.** The *per-capita* growth rate of strain A (A-B) or strain C (C-D) is shown against the density of strain A (A, C) or strain C (B, D), in the 7-7d fluctuation cycle. Each point corresponds to the *per-capita* growth rate, over the interval between two subsequent population measurements. The predictions from linear models (LM1, straight dashed lines and dark gray 95% CI) and generalized additive models (GAM1, continuous lines and light gray 95% CI) are also shown, pooling estimates from 1000 resimulated datasets to account for uncertainty in *r* and *N* (the competitor density was set at the median value of the observed densities). Here, none control mixtures satisfied the population density condition on both strains.

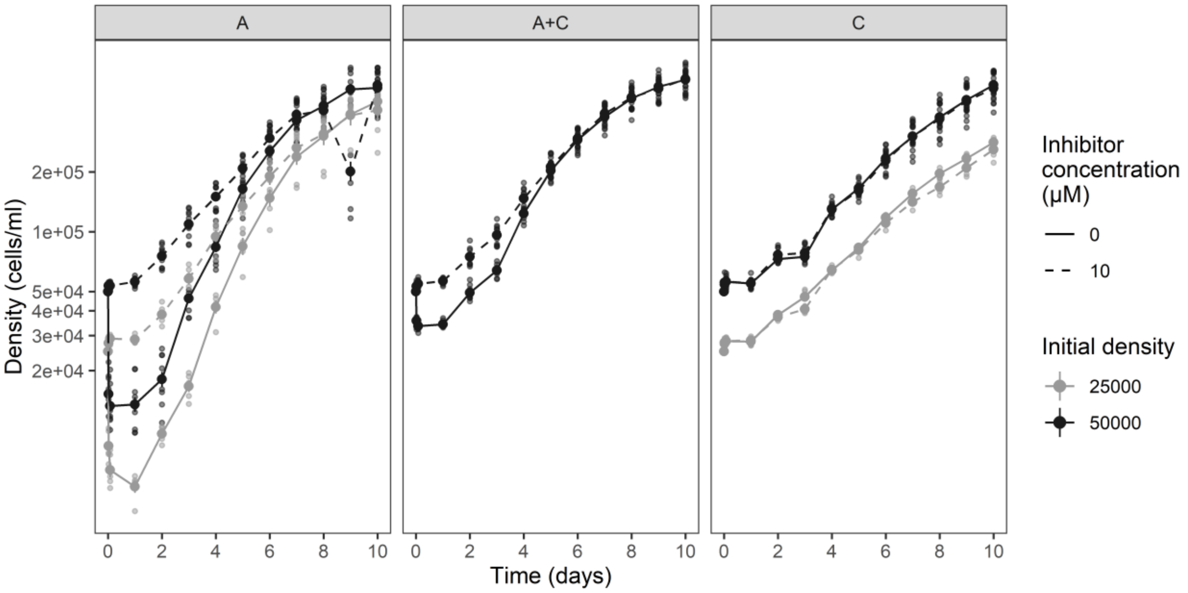


**Figure S7. Effect of PCD inhibitor treatment on demographic response after one salinity rise**. Populations densities of monocultures and mixtures (panels) are shown against time for different inhibitor treatments (line type), starting at different initial densities (colours). We computed the mean and standard error (not visible) on the cytometer counts over the isogenic lines (10 replicates for $N_0$= 50,000 cells/ml, 5 replicates for $N_0$ = 25,000 cells/ml).

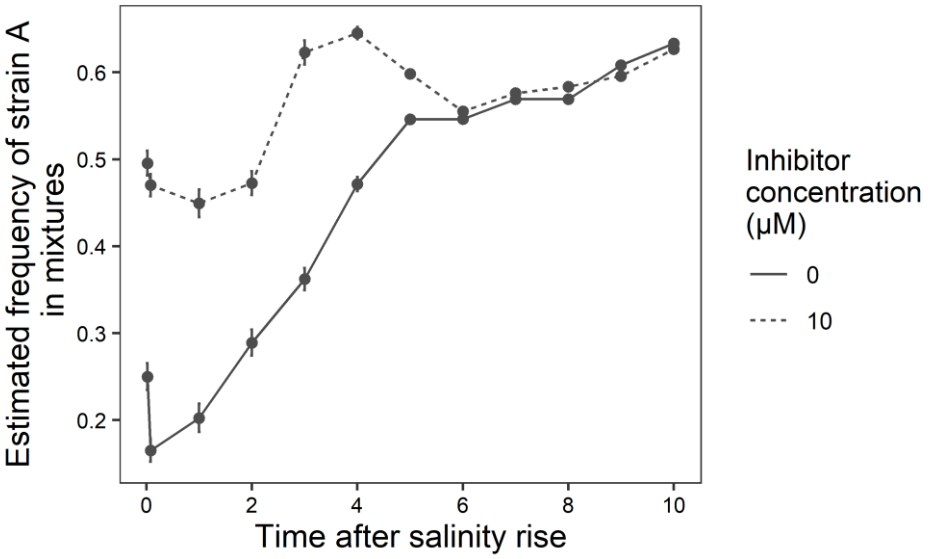


**Figure S8. Effect of PCD inhibitor treatment on strain A frequency in mixtures after one salinity rise**. The proportion of strain A in mixed populations, as estimated from the cytometric traits of cells, is shown against time for PCD inhibitor doses (line type). Means and confidence interval at 95% were obtained on the logit scale by inverse-variance weighting over the 10 population lines, to account for the variable uncertainty of estimates among biological replicates (isogenic lines of each strain), and then combined with the delta method for the back-transformation on the arithmetic scale.

#### Supplementary references


1. Leung C, Rescan M, Grulois D, Chevin LM. 2020 Reduced phenotypic plasticity evolves in less predictable environments. *Ecol. Lett.* **23**, 1664–1672. (doi:10.1111/ele.13598)

2. Leung C, Grulois D, Chevin L-M. 2022 Plasticity across levels: relating epigenomic, transcriptomic, and phenotypic responses to osmotic stress in a halotolerant microalga. *Mol. Ecol.*

3. Zeballos N, Grulois D, Leung C, Chevin L-M. 2023 Acceptable loss: Fitness consequences of salinity-induced cell death in a halotolerant microalga. *Am. Nat.* **201**. (doi:https://doi.org/10.1086/724417)

4. Bulmer MG. 1979 Principles of statistics. Courier Corporation.

5. Strimmer K. 2019 Statistical Methods: Likelihood, Bayes and Regression. , 213. See https://strimmerlab.github.io/publications/lecture-notes/MATH20802/maximum-likelihood-estimation.html (accessed on 28 May 2024).

6. Borenstein M, Hedges L V, Higgins JPT, Rothstein HR. 2021 *Introduction to meta-analysis*. John Wiley \& Sons.

7. Marin-Martinez F, Sánchez-Meca J. 2010 Weighting by inverse variance or by sample size in random-effects meta-analysis. *Educ. Psychol. Meas.* **70**, 56–73.

8. Ver Hoef JM. 2012 Who invented the delta method? *Am. Stat.* **66**, 124–127. (doi:10.1080/00031305.2012.687494)

9. Rescan M, Grulois D, Ortega-Aboud E, Chevin LM. 2020 Phenotypic memory drives population growth and extinction risk in a noisy environment. *Nat. Ecol. Evol.* **4**, 193–201. (doi:10.1038/s41559-019-1089-6)

10. Rubin DB. 1987 *Multiple Imputation for Nonresponse in Surveys*. John Wiley.

11. Miles A. 2016 Obtaining Predictions from Models Fit to Multiply Imputed Data. *Sociol. Methods Res.* **45**, 175–185. (doi:10.1177/0049124115610345)

12. Kot M. 2001 *Elements of Mathematical Ecology*. Cambridge University Press.
